# Reversable Acute Sedation Response of Phosphorothioate Antisense Oligonucleotides Following Local Delivery to the Central Nervous System

**DOI:** 10.1101/2025.02.13.638136

**Authors:** Jacqueline G. O’Rourke, Gemma Bachmann, Curt Mazur, Keming Zhou, Oleksandr Platoshyn, Mariana Bravo Hernandez, Stephanie Klein, Jonathon Nguyen, Sebastien Burel, Christine Hoffmaster, Tom Zanardi, Paymaan Jafar-nejad, Martin Marsala, Scott P. Henry, Eric E. Swayze, Berit Powers, Holly B. Kordasiewicz

**Affiliations:** Ionis Pharmaceuticals, Inc., Carlsbad, CA 92008, USA; University of California San Diego School of Medicine, Department of Anesthesiology, La Jolla, CA 92093, USA

**Author notes:** Corresponding Author: Holly Kordasiewicz, Ionis Pharmaceuticals, Inc., Carlsbad, CA.

## Abstract

Antisense oligonucleotides (ASOs) locally delivered to the central nervous system (CNS) are being approved as therapies for neurological diseases. After intrathecal injection of some ASOs, transient toxicities have been reported, but considerable inconsistencies remain in classifying them and their underlying mechanisms. Here, we characterize an acute sedation response that can include loss of lower spinal reflexes, hypoactivity, paresis, sedation and ataxia, peaking ∼3 hours post-intrathecal injection of some phosphorothioate ASOs and reversing by 24 hours with no sequelae. Acute sedation is distinct from acute activation, which is hyperactivity and muscle cramping that occurs immediately after administering oligonucleotides. Acute sedation translates across species from rodents to non-human primates and is sequence-, dose-, and chemistry-dependent. Acute sedation can be mitigated by strategic placement of phosphorothioate backbone linkages in ASOs and by avoiding G-rich sequences. The acute sedation response can be modeled in primary neural cultures, with good predictability of *in vivo* response. Mechanistically, we demonstrate that acute sedation is caused by high extracellular ASO concentrations inhibiting synaptic transmission, which reverses as ASO is cleared from the extracellular space and taken up into cells. Our results provide a comprehensive framework for quantifying and mitigating acute sedation caused by some phosphorothioate ASOs.

**GRAPHICAL ABSTRACT:** 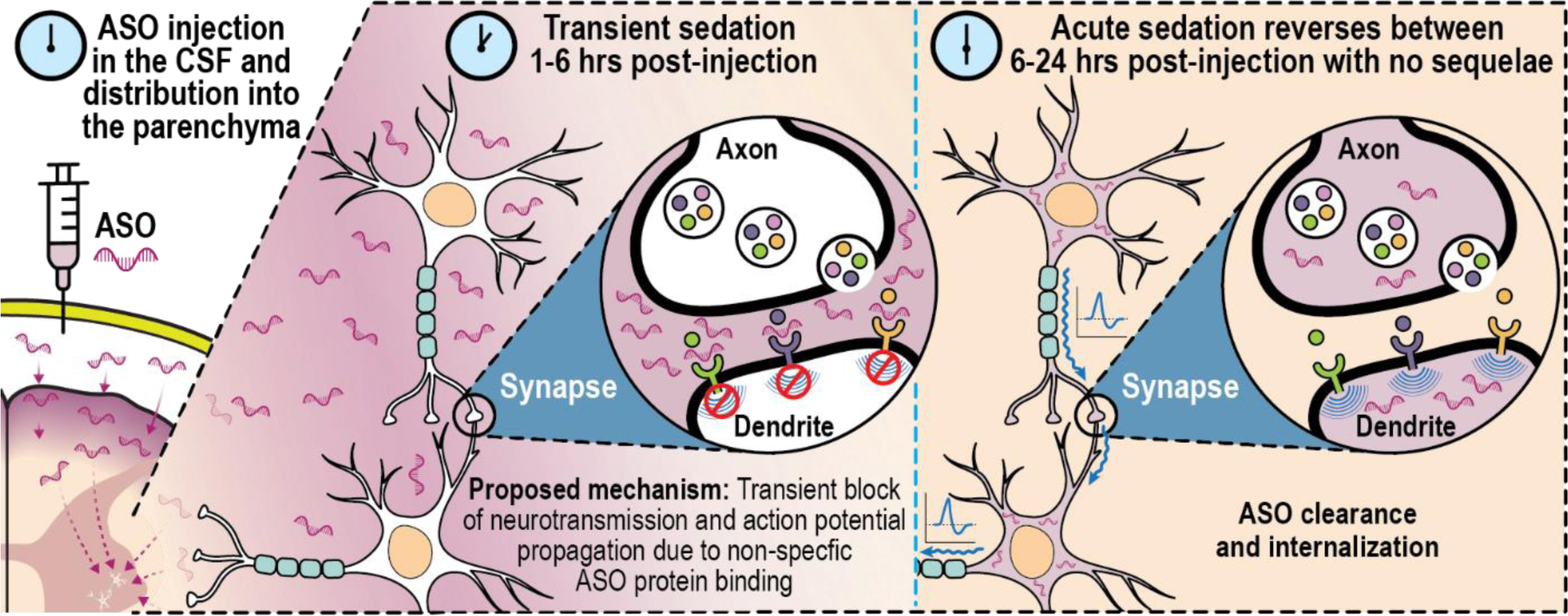

## INTRODUCTION

Antisense oligonucleotides (ASOs) are synthetic nucleotides 12-30 bases in length that modulate target RNAs [1]. Two locally delivered ASOs have been approved for the treatment of neurological disorders, nusinersen (Spinraza) for the treatment of spinal muscular atrophy [2, 3] and tofersen (Qalsody) for the treatment of SOD1-mediated amyotrophic lateral sclerosis (ALS) [4]. Because ASOs modulate RNA, they hold tremendous promise for the treatment of a broad range of CNS disorders. Indeed, many locally delivered ASO therapies are now in clinical development [5-10].

Because ASOs are relatively large, highly charged molecules, they do not cross the blood-brain-barrier and are instead administered by local intrathecal (IT) delivery into the cerebrospinal fluid (CSF) [11]. This results in high local concentrations of ASO in the intrathecal space surrounding the spinal cord and nerve roots within the minutes to hours after dosing, before the ASO distributes broadly throughout the CNS parenchyma [12, 13]. Nonspecific interactions between phosphorothioate oligonucleotides and proteins are a well-known property associated with systemic administration of ASOs [14]. Taken together, it is not unexpected there have been previous reports of transient responses that occur in the minutes to hours after administration of ASOs into the CNS of rodents and non-human primates (NHP). These include behaviors indicative of nervous system depression or sedation including decreased spinal reflexes, motor activity, and muscle strength, as well as ataxia, decreased consciousness, slowed breathing, and abnormal posture [15-18]. These reports have also described responses indicative of nervous system activation such as increased locomotor activity, nystagmus, stereotypy, muscle tremors, spasticity, convulsions and seizures. Often all types of observed acute symptoms, whether indicative of CNS sedation or activation, are captured together with the data presented as one composite acute response score. Furthermore, the kinetics of the development and resolution of different responses are often ignored or not reported. This approach makes the overall behavioral findings and any investigations into causal mechanisms difficult to interpret. As the field continues to hone the process of identifying safe and efficacious ASOs for use in the clinic, there is a need to carefully define potential liabilities, develop tools to translate these findings across species, and to identify, eliminate, or modify poorly tolerated compounds. This is our aim.

Here, we characterize and classify a sequence-, dose- and chemistry modification-dependent acute sedation response following ASO administration into the CNS that translates across preclinical species – mice, rats, and NHP. This response is due to high extracellular concentrations of ASO, which transiently inhibit synaptic transmission. We demonstrate this is distinct mechanistically from the acute activation response, described for both siRNAs and ASO [15-18]. We demonstrate how the acute sedation effect can be captured in a neurobehavioral scale, how this translates across species, and how it can be modeled in an *in vitro* screening assay.

## MATERIALS AND METHODS

### ASO preparation

Lyophilized ASOs (Supplementary Table S1 and S2) were dissolved in sterile phosphate-buffered saline (PBS) without Ca^2+^ or Mg^2+^ for all rodent studies, unless otherwise noted. For dosing solutions containing supplemental concentrations of Ca^2+^ or Mg^2+^, CaCl_2_ and MgCl_2_ stock solutions were formulated in aCSF with the addition of divalent cations (Ca^2+^ or Mg^2+^). The aCSF used already contained 2.2 mM divalent cations, which was subtracted out of the final concentration of Ca^2+^ or Mg^2+^ added. ASO was quantified by UV spectrometry and was sterilized by passage through a 0.2-µm sterile before dosing.

### Intracerebroventricular (ICV) administration in mice

All protocols met ethical standards for animal experimentation and were approved by the Institutional Animal Care and Use Committee of Ionis Pharmaceuticals. Female C57BL/6J mice were obtained from the Jackson Laboratory (stock # 000664; Bar Harbor, ME). ICV injections were performed as previously described [12]. Briefly, mice were anesthetized with isoflurane and placed in a stereotaxic frame, and an incision was made in the scalp. A Hamilton microsyringe was placed at 0.3 mm anterior and 1.0 mm lateral to bregma and was lowered 3.0 mm into the lateral ventricle. A 10 μl volume of ASO solution was injected at a rate of 1 μl/s and the needle held in place for 3 min before being withdrawn and the incision closed. The mice were returned to their home cages to recover from anesthesia.

### Intrathecal (IT) administration in rats

IT administration of ASO in rats was performed under a protocol approved by the Institutional Animal Care and Use Committee of Ionis Pharmaceuticals. Male Sprague-Dawley rats (n = 4/group) with body weights between 0.25 and 0.4 kg were obtained from Harlan laboratories (stock # 213M, Livermore, CA). Catheterization of the lumbar intrathecal space was performed as described previously [19]. ASO or vehicle was delivered over 30 seconds in a 30 μl volume followed by 40 μl of vehicle. The animals were returned to their home cages for recovery from anesthesia.

### Intrathecal (IT) administration of ASO in NHP

NHP studies were performed at Northern Biomedical Research, Inc., Spring Lake, MI and were approved by their Institutional Animal Care and Use Committee. As previously described [12], adult male or female cynomolgus monkeys were anesthetized and 1 ml aCSF or ASO dosing solution was administered via percutaneous IT injection over 1 minute using a spinal needle at the lumbar level (target L4/L5 space). Animals were dosed in lateral recumbency position and remained in a prone position for at least 15 min following dosing.

### Acute sedation scoring following IT administration in rodents

Animals are lifted out of their home cages and observed on a flat surface 3 hours after ASO IT administration. They are assessed according to the scale in Supplementary Figure S1A and an acute sedation score is assign and referred to as the 3 hour acute sedation score (3h aS score (0-7). A score of 0 is bright, alert, responsive. The end of the tail is gently lifted to hip level and let go. If it hits the table without muscle tone it scores a 1. If the hind end is drooping or abnormal gait is present in hindlimbs, a score of 2 is given. If the hindlimbs are unable to support weight, but hind paws can still move, it scores a 3. If the hindlimbs are not moving at all, it scores a 4. If the animal is unable to support their front half with forelimbs, it scores a 5. If the animal cannot move its forelimbs, and there is any movement at all, including breathing, it scores a 6. Death is scored as a 7.

### Acute sedation scoring following ICV administration in rodents

Animals are observed in their home cages 3 hours after ASO ICV injection. They are assessed according to the scale in Supplementary Figure S1B and an acute sedation score is assign and referred to as the 3 hour acute sedation score (3h aS score (0-7)). A score of 0 is bright, alert, and responsive. A score of 1 is mild gait impairment (i.e. wobbly walking). A score of 2 is obvious impaired gait and/or a hunched posture. If the animal is lying on its side and is not ambulating in the home cage, it is lifted out and placed on a flat surface. If it moves away from the experimenter after lifting, it scores a 3. If the animal does not move forward after lifting, but has an upright posture, it scores a 4. If the animal is lying on its side and does not move after lift, but responds to tail pinch, it scores a 5. If it does not respond to pinch, but is breathing, it scores a 6. Death is scored as a 7.

### Acute spinal reflex scoring following IT administration in NHP

Lower spinal reflexes and general motor function are assessed 2-4 hours after IT administration of ASO as follows. Lack of response in each area is given a score of 1 (movement = 0, lack of movement = 1) and a cumulative score is assigned for each animal. Tail Reflex: When the tail is touched with a sharp object, the tail should move to avoid the stimulus. Cutaneous Reflex: The reaction to a series of needle pricks along each side of the abdomen was recorded. Sensory Foot Reflex: The foot was stimulated with a sharp object and an avoidance response was observed. Proprioceptive Reflex: Each foot was inverted and the ability to right the foot was noted. Knee Jerk Reflex: The presence or absence of a quick extension of a hind leg was noted following the tapping of the patellar ligament with a reflex hammer.

### Grip strength and open field assessments of motor function in mice

For all behavioral tests, mice were acclimated to the testing room for a minimum of 60 minutes prior to each round of testing. Grip strength: The Chatillon-Ametek Digital Force Gauge, DFE II (Columbus Instruments, Columbus, OH) was used to determine the strength exerted by the hindlimbs of each animal in response to a constant downward force. Data presented as average of 4 trials as grip force (Gf) normalized to body weight (Gf/g). Open field: Kinder Scientific Open Field Arenas with 16 x 16 beams located 1 inch apart were used for this test. Mice were placed individually into the center of the arena and total activity was measured as the number of beam breaks over 300 seconds.

### *In vivo* electrophysiological recordings

Rats were instrumented for stimulation in the motor cortex or tibial nerve and recordings in the thoracic spinal cord, lumbar spinal cord or peripheral muscle. For each test of the electrophysiological recordings the animals were dosed via IT administration with vehicle to measure a baseline recording, then with ASOs. Electrophysiological recordings were taken 2 and 24 hours after dosing with the ASO.

### Myogenic motor-evoked potentials recording (MMEP)

MMEPs were measured as previously [20]. Briefly, animals were anesthetized with propofol (50-100 mg/kg, i.p) and two 30-G stainless steel stimulating electrodes were placed subcutaneously overlying the left and right motor cortex. Motor-evoked potentials were elicited by transcranial electrical stimulation with a pulse duration of 1 ms at 7.5 mA using a DS3 constant current isolated stimulator (Digitimer). Responses were recorded from the gastrocnemius muscle using 30-G platinum transcutaneous needle electrodes (distance between recording electrodes ∼1 cm; Grass Technologies, Astro-Med).

### Spinal motor-evoked potentials recording (SMEP)

Spinal cord surface motor-evoked potentials were recorded from the dorsal surface of the lower thoracic (Th12) spinal cord as previously [21]. Under isoflurane anesthesia [2.0–2.5% (vol/vol) maintenance; in room air], animals were mounted into a stereotaxic frame. A dental drill was used to perform a laminectomy of T11 vertebra exposing the T12 spinal segment. Two 30-G stainless steel stimulating electrodes were placed subcutaneously overlying the left and right motor cortex.

Spinal cord motor-evoked potentials were elicited by transcranial electrical stimulation with a pulse duration of 0.2 ms at 7.0 mA using a DS3 constant current isolated stimulator (Digitimer). Evoked responses were recorded by a pair of flexible silver-ball electrodes placed on the dura surface of the exposed T12 spinal segment (distance between recording electrodes∼0.5 cm). A reference silver-chloride disk electrode was placed subcutaneously on the contralateral side of the recording. After electrode placement, animals were injected with ketamine propofol (50-100 mg/kg, i.p) and isoflurane anesthesia was discontinued.

### Spinal somato-sensory evoked potentials recording (SSEP)

Spinal cord surface somato-sensory evoked potentials were recorded from the dorsal surface of the lumbar (L2-L5) spinal cord as previously [22]. Under isoflurane anesthesia [2.0–2.5% (vol/vol) maintenance; in room air], animals were mounted into a stereotaxic frame. A dental drill was used to perform a laminectomy of lumbar vertebra exposing the L2-L5 spinal segment. Two 30-G stainless steel stimulating electrodes were transcutaneously inserted into the surroundings of the tibial nerve (distance between stimulation electrodes ∼1 cm; Grass Technologies, Astro-Med). Spinal cord surface somato-sensory evoked potentials were elicited by transcranial electrical stimulation with a pulse duration of 0.2 ms at 6.0 mA using a DS3 constant current isolated stimulator (Digitimer). Evoked responses were recorded by a pair of flexible silver-ball electrodes placed on the dura surface of the exposed L2-L5 spinal segment (distance between recording electrodes ∼0.5 cm). A reference silver-chloride disk electrode was placed subcutaneously on the contralateral side of the recording. After electrode placement, animals were injected with propofol (50-100 mg/kg, i.p.) and isoflurane anesthesia was discontinued.

### Hoffman reflex (HR; a proprioceptive reflex)

HR was recorded as previously [23]. Under isoflurane anesthesia, the right hindlimb of an animal was secured, and a pair of stimulating needle electrodes was transcutaneously inserted into the surroundings of the tibial nerve. For recording, a pair of 30-G platinum needle electrodes were inserted into the interosseous muscles between the fourth and the fifth, or the first and the second metatarsal muscles of the right hind paw, and a grounding electrode was placed into the tail. The tibial nerve was stimulated using square pulses with increasing stimulus intensity (0.1–10 mA in 0.5-mA increments, 0.1 Hz, 0.2 ms; DS3 constant current isolated stimulator, Digitimer, Hertfordshire, UK). We used these data to determine the intensity necessary to obtain a maximal M and H response. All recording (from MMEP, SMEP, SSEP and HR) electrodes were connected to an active head stage (3110W Head stage; Warner Instruments) and signal-amplified (x100 times) using DP-311 differential amplifier (Warner Instruments). Amplified signal was acquired by the PowerLab 8/30 data-acquisition system (ADInstruments) at sampling frequency of 20 kHz, digitized and stored in PC for analysis.

### Tissue harvesting

Mice and rats were euthanized 1 hour to 2 weeks after the ASO injection as indicated by the study. Frozen brain and spinal cord tissues were harvested for the determination of ASO concentration and target mRNA expression as described below. Tissues were also immersion fixed in 10% buffered formalin solution for histological processing as described below.

### Quantitative real-time PCR

RNA was extracted and qRT-PCR performed as previously described [12] using gene-specific primers (IDT technologies, Coralville, IA). For ASO treated mice the expression level of *Malat1* was normalized to that of *Gapdh* and this was further normalized to the level in vehicle treated animals. Primer/probes sequences: *Malat1* forward – AGGCGGGCAGCTAAGGA; *Malat1* reverse – CCCCACTGTAGCATCACATCA; *Malat1* probe – TTCCTCTGCCGGTCCCTCGAAAG; *Gapdh* forward – GGCAAATTCAACGGCACAGT; *Gapdh* reverse – GGGTCTCGCTCCTGGAAGAT; *Gapdh* probe – AAGGCCGAGAATGGGAAGCTTGTCATC.

### Quantification of ASO tissue levels

ASO6 and ASO6_PO were quantified in rat CNS tissue a previously described [24]. Briefly, samples were weighed, homogenized, and ASO was extracted from the tissue via liquid–liquid extraction. The aqueous layer was processed via solid phase extraction with a Strata X plate. Eluates were passed through a protein precipitation plate before drying down under nitrogen at 50°C. Dried samples were reconstituted in 140 μl water containing 100 μM EDTA and were run on an Agilent 6130B single quadrupole LCMS to quantify ASO concentration.

### Generation of mouse primary cortical cultures and multielectrode array (MEA) recordings

Mouse cortical cells were isolated from C57BL/6 embryos on day 16 and plated in MEA plates pre-coated with Poly-D lysine at 150K to 200K per well and cultured in Neurobasal media. Neurons were cultured for 2 weeks on 24 well glass bottom plates from Multi Channel Systems containing 12-electrodes before MEA recordings and ASO treatments. For MEA recordings, MEA plates were allowed to acclimate before recording a 10 min baseline for all wells. Then primary neurons were treated with ASO at various concentrations (from 5-150 µM) and recorded for an additional 10 min. For washout experiments, after recording the ASO treatment phase, wells were flushed with media 2 times and then recorded for neuronal activity on the MEA. Percent inhibition was calculated as 100% X (1-Spike Rate (ASO)/Spike Rate (baseline)) and if Spike Rate ASO > Spike Rate baseline percent inhibition was recorded as 0.

### Histology

Rat brains were processed as described previously [12]. Briefly, they were bisected sagittally at the midline, immersion fixed in 10% Neutral Buffered Formalin (Statlab, 28600-1) for 72 h, transferred to 70% ethanol, processed into Paraplast Plus paraffin (Leica, 39602004) overnight on a Sakura Tissue Tek tissue processor, and embedded in paraffin. Tissue sections were cut at 4 μm thick and mounted on positively charged slides (Millennia 1000, Statlab, M1000W), then air dried overnight and dried at 60°C in a slide drying oven for 1 h prior to staining.

### ASO immunohistochemistry

Slides were stained with Rabbit polyclonal ASO (Ionis) antibody on a Ventana Ultra staining system. ASO slides were treated enzymatically with Trypsin (Sigma, T8003). The slides were then blocked with Endogenous Biotin Blocking Kit (Ventana, 760-050) and Normal Goat Serum (Jackson Immuno Labs, 005-000-121). The primary antibody was diluted with Discovery Antibody Diluent (Ventana, 760-108) and incubated for 1 hour at 37°c. The antibodies were detected with Biotin labeled Goat Anti Rabbit secondary antibody (Jackson Immuno Labs, 111-005-003). The secondary antibody was labeled with DABMap Kit (Ventana, 760-124). Note: ASO antibody staining level is dependent on ASO sequence and phosphorothioate (PS) content in the backbone.

Images were scanned on a Hamamatsu S360 scanner at 20X resolution.

### Data analysis

All data are shown as mean ± SD, and all analyses were conducted using Prism software (GraphPad Software, San Diego, CA). Grip Strength and Open Field statistical analysis determined by two-way ANOVA. Acute Sedation Score graphs were determined by nonparametric tests such as Kruskal-Wallis test with Dunn’s multiple comparison or Mann-Whitney and stated in the figure legends. Correlation graphs determined by Spearman r test. ED50 determined by non-linear regression with normalized response and variable slope (Motulsky).

## RESULTS

### Acute sedation is dose- and concentration-dependent following local CNS ASO administration

We observed that certain ASOs caused transient loss of muscle strength and paresis/paralysis that presented within the first few hours after local intrathecal (IT) administration and generally resolved between 6-24 hours post-injection. To characterize the dose dependence of the response, we administered an ASO known to cause the response, ASO1, at 100, 300, 700, or 1000 µg via IT injection in rats and assessed the motor behavior at 1-8 hours and at 24 hours post-administration (Figure 1A). Motor deficits were present in just a few regions in the posterior aspect (i.e. tail, posterior posture, hind limbs) at low doses. At higher doses, additional regions were affected in a progressively anterior pattern (i.e. full hindlimb paralysis, then anterior posture, then forepaws). By 1-hour post-dose paresis was present in posterior regions, then moved further anteriorly to include more regions and peaked between 3-4 hours. Paresis/paralysis began to wane between 5-8 hours, wearing off in an anterior to posterior pattern.

**Figure 1.**
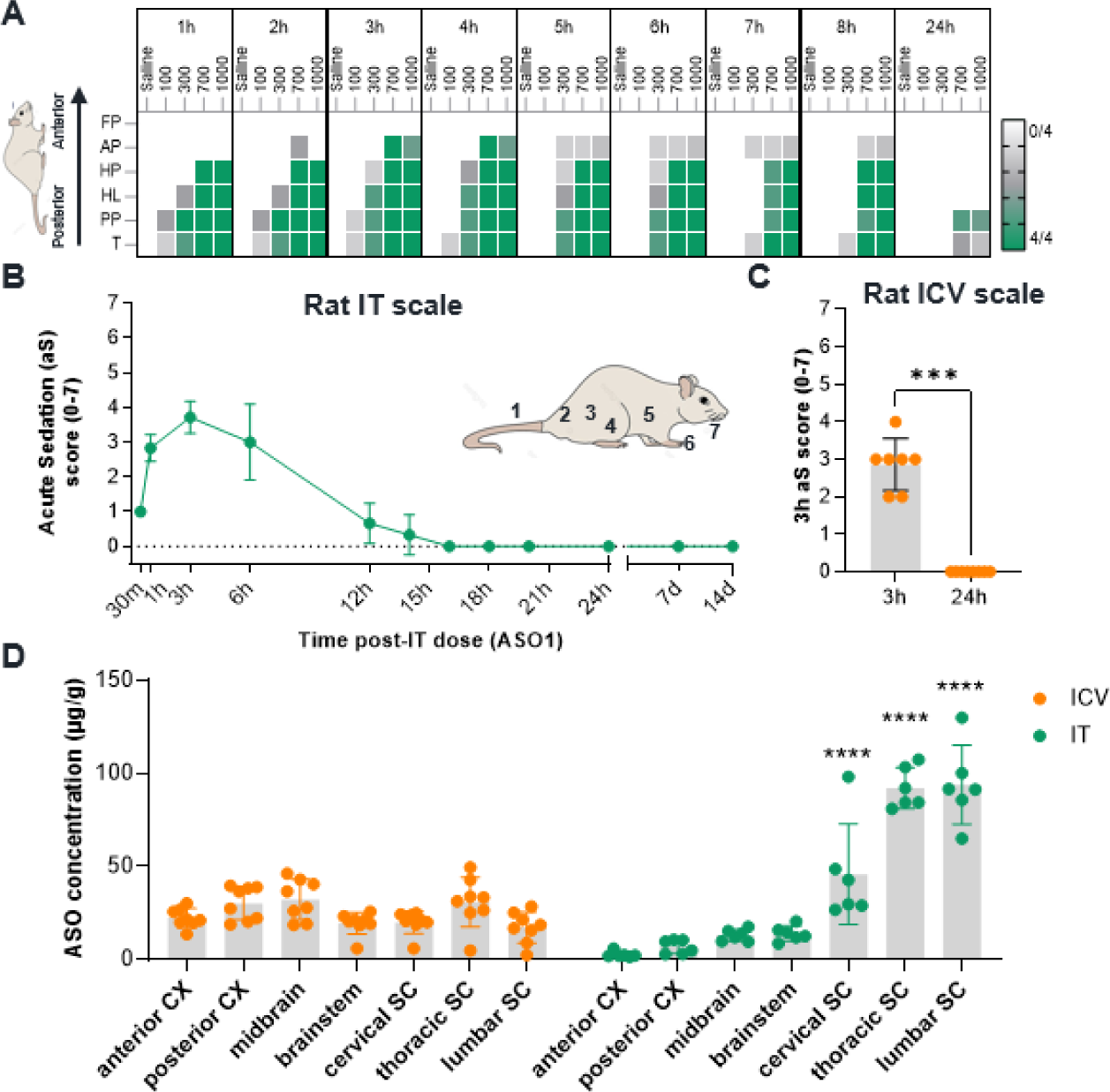
ASO-mediated acute sedation response is dose-dependent and driven by level of tissue exposure in different CNS regions. (**A**) Wild-type rats (n=4) were dosed with vehicle (saline), or 100 µg, 300 µg, 700 µg, or 1000 µg of ASO1 via IT administration and assessed for motor function at 1-8 hours (h) and at 24h post-dose. The heatmap shows the number of rats/group that exhibited paresis/paralysis in the designated regions over time (white = 0/4; green = 4/4). The response started in the posterior region and progress to anterior regions over the first 3-4h post-dose, then resolved between 5-24h in an anterior to posterior direction. Lower doses caused paresis in posterior regions only while higher doses also affected anterior regions. (**B**) Time course of acute sedation (aS) as scored on the scale developed for IT administration in rodents. Rats (n=3) received 700 µg of ASO1 and exhibited a low acute sedation score of 1 at 30 minutes (min) that increased to a peak of 3-4 at 3h post-injection. Acute sedation scores returned to 0 by 16h post-dose and continued at 0 out to 14 days (d) post-dose, indicating no sequelae. (**C**) Acute sedation as scored on the scale developed for ICV bolus injections in rats shows a similar level of acute sedation at 3h post-ICV delivery of 700 µg of ASO1, however the peak score of ∼3 indicates general sedation and lack of spontaneous ambulation without stimulus (**D**) Comparison of ASO tissue concentrations across CNS regions following ICV (orange) or IT (green) administration of 700 µg ASO1. Similar levels of ASO concentration across tissue 3h post-ICV injection whereas the higher ASO concentrations in the spinal cord vs. the brain regions with lumbar having the highest concentration following IT administration. Nonparametric test using Mann-Whitney test for statistical analysis in **C**. One-way ANOVA multiple comparisons post-hoc test were used to compare tissue concentration across different routes of administration in different tissues in **D**. *P < 0.05, **P < 0.01, ***P < 0.001, ****P < 0.0001. T = Tail; PP = posterior posture; HL = hindlimbs; HP = hind paws, AP = anterior posture, FP = forepaws, CX = cortex, SC = spinal cord.

We adopted the above observations into a scale to efficiently quantify the level of acute sedation after IT injection of ASO in rodents (Supplementary Figure S1A). The scale gives one point for the presence of paresis/paralysis in the seven regions, as depicted in Figure 1B. To quantify the response across time, we repeated the experiment with 700 µg of ASO1 delivered via IT injection in rats and assessed them from 30 minutes to 24 hours post-dose, and 1- and 2-weeks post-dose. At 30 minutes, all animals exhibited a score of 1 (tail paralysis; Fig 1B). By 3 hours, peak scores of 4 were reached. By 16-24 hours, scores of 0 (indicating bright, alert, responsive) were reached in all animals indicating full reversal of acute sedation. Scores of zero were maintained out to 2 weeks post-injection confirming acute sedation is transient with no long-term sequelae. For subsequent studies, we focused on the peak of the acute sedation response at 3 hours post-dosing.

We hypothesized that since acute sedation presented in a dose dependent, posterior-to-anterior pattern following IT administration, the acute sedation response was likely due to a posterior-to-anterior ASO concentration gradient in CNS tissues in the hours after IT dosing. This is consistent with the previously published gradient that occurs at dosing, and the time it takes for ASO to distribute up the neuraxis [13]. To assess this, we delivered ASO to the CSF via two routes, IT and intracerebroventricular (ICV), to vary the distribution of ASO across different brain and spinal cord regions in the hours after dosing. We then evaluated behavior at 3 hours post-dosing and quantified tissue concentrations of ASO. As before, the IT-delivered ASO1 caused an acute sedation response involving loss of lower limb mobility (Figure 1C). The response presented differently following dosing of the same ASO delivered at the same dose level, but via ICV injection. Here, ASO1 caused a general sedation more consistent with inhibition of brain function than spinal cord function. We translated these observations into a similar 7-point scoring system for quantification of acute sedation response after ICV in rodents, to allow for quantitative comparisons across delivery routes (Figure 1C, Supplemental Figure S1B). Tissue concentrations confirmed that at early timepoints post-dosing, ASO delivered via IT and ICV distribute differently, and the CNS regions with the highest concentrations are most affected by the acute sedation response (Figure 1D).

### Acute sedation response is sequence-dependent and reversible

In mice, ASO1 delivered by ICV injection, caused a similar response over a similar time course to that in rats following ICV injection (Supplementary Figure S2B). IT injection in mice is difficult to perform with consistent delivery up the neuraxis, thus all mice in these studies were assessed by ICV delivery. To determine the sequence dependence of the acute sedation response and to more deeply characterize the reversibility, we selected a panel of seven ASOs with different nucleotide sequences but identical chemistries, fully phosphorothioate (PS) modified 20-mer oligonucleotides with a 10-base deoxynucleotide gap flanked on each end with 5 2’-O-Methoxyethyl (MOE) modified nucleotides (ASOs 2-8; Supplementary Table S1). ASOs (100 µg) or vehicle were administered via ICV injection to mice. Based on the time course of acute sedation peak and resolution observed in rats and mice (Figure 1A-C; Supplementary Figure S2A), we assessed the acute sedation response at 3 hours and 24 hours post-ICV injection. The acute sedation response was sequence-dependent (Figure 2A, left panel). Mean acute sedation scores were significantly different and ranged from 0-4. Acute sedation in all the mice was resolved by 24h post-injection (Figure 2A, right panel). To better characterize the reversal, we used well-established sensitive locomotor behavior assays, grip strength [25] and open field [26]. At 3 hours post-dosing, for ASOs with a high acute sedation response there was a significant reduction in grip-strength (Figure 2B), and locomotion in the open-field (Figure 2C), and this reversed by 24 hours post-dosing (Figure 2A-B). Some sequences exhibited no change in any behavior (ASO2 and 3). The acute sedation scores significantly correlated with both grip strength and locomotion at 3 hours post-dosing (Grip strength, r = -0.7611 and open-field r, = -0.8463; Supplementary Figure S2C).

**Figure 2.**
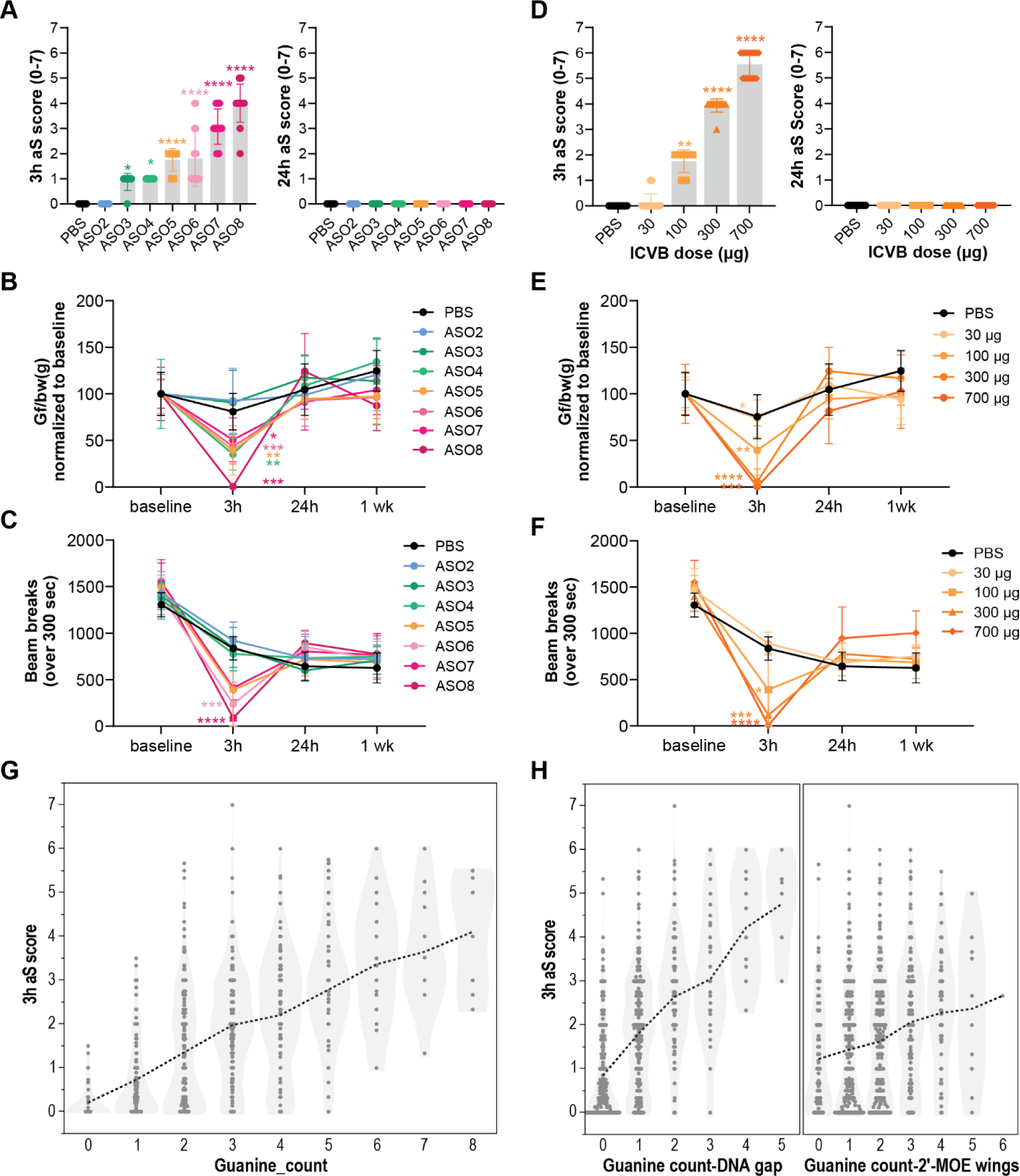
ASO-mediated acute sedation responses are transient, sequence dependent with no sequalae. (**A**) Mice (n=16) ICV (ICV) injected with ASO2-8 (100 µg) were scored at 3h and 24h post-injection using the acute sedation (aS) scale (0-7). ASO2-8 showed a range of acute sedation scores (0 to 4) at 3h and the responses were reversed by 24h. (**B**) Mice (n=8) received 100 µg of full-PS backbone MOE gapmer ASOs (ASO2-8) via ICV injection. They were assessed for hindlimb Grip Strength (GS) at pre-dose (baseline) and at 3 hours (h), 24h and 1-week (wk) post-dose. ASOs 4-8 significantly suppressed GS at 3h post-dose to differing degrees compared to baseline. GS recovered by 24h and 1 wk-post dose in all cases. (**C**) Mice (n=8) received 100 µg of ASO2-8 via ICV injection and the total activity was measured by the number of beam breaks in 300 seconds in an Open Field (OF) assay at pre-dose (baseline) and at 3h, 24h and 1-wk post-dose. (**D**) Acute sedation scores in mice following ICV injection of ASO5 at increasing doses (30 µg, 100 µg, 300 µg, and 700 µg) and scored at 3h and 24h post-dose. (**E**) Mice (n=8) received different dose levels of ASO5 (30 µg, 100 µg, 300 µg, or 700 µg) via ICV injection. They were assessed for hindlimb GS at 3h, 24h and 1-wk post-dose. A dose-dependent suppression of GS was observed. (**F**) Mice (n=8) received different dose levels of ASO5 (30 µg, 100 µg, 300 µg, or 700 µg) via ICV injection and total activity was assessed by OF at pre-dose (baseline) and at 3 hours, 24 hours and 1-week post-dose. Total activity was suppressed by ASOs 6 and 8 at 3h post-injection, with recovery by 24h. **G**) Total number of Guanines (Gs) in a given ASO sequence compared to the 3h acute sedation score following 700 µg ICV injection in mice. (**H**) Number of Gs present in either the DNA gap (left panel, bases 6-15) or 2’-MOE wings (right panel, bases 1-5 and 16-20) relative to the 3h acute sedation score following 700 µg ICV injection in mice. Each data point represents the mean and error bars represent standard deviations. **A** and **D** *P < 0.05, **P < 0.01, ***P < 0.001, ****P < 0.0001 using the nonparametric Kruskal-Wallis test and Dunn’s multiple comparisons post-hoc test, **B**, **C**, **E** and **F** using two-way ANOVA.

To further characterize the dose-dependency of acute sedation and reversibility (Figure 1A), we selected ASO5 which exhibited a moderate response at 100 µg and administered it in increasing dose levels from 30 µg up to 700 µg via ICV injection in mice. At the top dose, ASO5 resulted in an acute sedation score of 6, which is the highest score an animal could achieve and survive (Figure 2D). Both grip strength (Figure 2E) and open field (Figure 2F) were significantly impaired at 3 hours, with mice treated with 700 µg of ASO5 exhibiting no locomotion in the open field (Supplementary Figure S2B). All impairments reversed by 24 hours. Taken together, different sequences of phosphorothioate ASOs have a different magnitude of acute sedation response that is dependent on dose level, ranging from none to severe acute sedation, and this response is reversable by 24 hours post-dosing.

### G-rich, C-poor sequences are acutely sedative

To better understand the sequence contributions to the acute sedation phenotype, we assessed the base content of the ASOs. ASO sequences containing an enrichment of guanine (G) nucleotides were previously associated with suppression of synchronous neuronal activity in an *in vitro* model of ASO toxicity [16], while conversely adenine (A)-rich sequences were associated with maintenance of neuronal activity. G-rich sequences at the 3’ end of the ASO were found to severely suppress neuronal activity. We analyzed the nucleotide content and placement in ∼1800 ASOs compared to their *in vivo* acute sedation scores and found the severity of acute sedation to be positively associated with enrichment for G nucleotides (Figure 2G), whether in the DNA gap or the 2’-MOE wings (Figure 2H). Furthermore, we found high cytosine (C) content and adenosine (A) content was associated with a trend in low acute sedation scores, while thymine (T)-rich sequences bore no specific effect on severity of acute sedation (Supplementary Figure S2D). This data indicates sequence features can profoundly affect acute sedation with G and C having opposing effects.

### Acute sedation response translates across species

To determine if the acute sedation response translates across species, we correlated the response of over 1000 ASOs that were tested in both mice and rats, animals were scored at 3 hours post-dose following treatment with a 700 µg ICV injection in mice and a 3000 µg IT administration in rats. We found acute sedation scores were highly correlated (R^2^= 0.414, Figure 3A), indicating translatability across species. The response does not scale by CSF volume, with rats receiving an IT ASO generally being more sensitive than mice with an ICV delivered ASO, potentially due to the different routes of administration.

**Figure 3.**
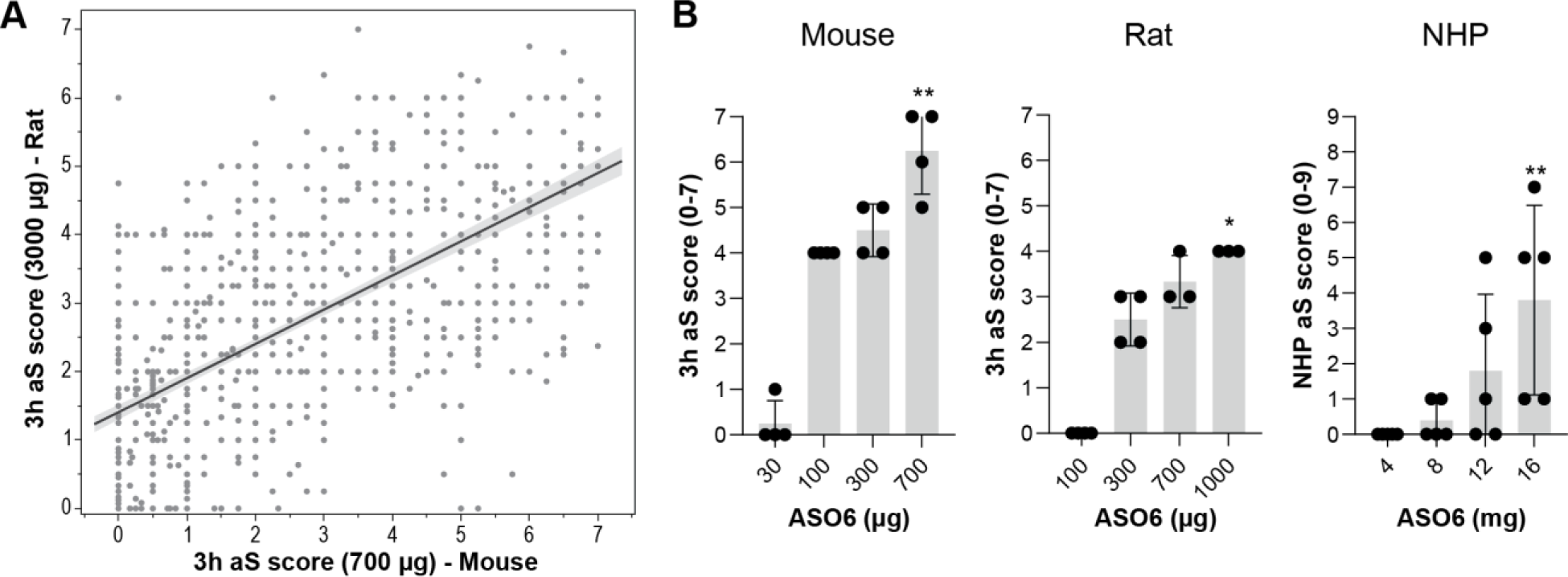
Acute ASO sedation responses translate across preclinical species and are dose dependent. (**A**) Correlation plots of acute sedation (aS) scores for different ASOs 3 hours (h) post-injection in rat (3000 µg IT) and mouse (700 µg ICV). (**B**) Dose-dependent aS scores observed with ASO6 in mice, rats, and NHPs. Each data point represents individual animals’ scores. Dose-dependent increases in incidence of sedation responses are observed. *P < 0.05, **P < 0.01, ***P < 0.001, ****P < 0.0001 using the nonparametric Kruskal-Wallis test and Dunn’s multiple comparisons post-hoc test in **B**.

To directly compare acute sedation across species, we administered ASO6 in dose response studies in mice, rats, and NHP and scored the acute sedation response 3-4 hours post dosing. As demonstrated previously (Figure 2), ASO6 again had a clear acute sedation response in mice following ICV injection (top dose mean score of 6.25, p < 0.01; Figure 3B), and in rats following IT (top dose mean scores of 4, p < 0.05; Figure 3B). Like IT in rats, acute sedation in NHP after IT administration presents as a loss of spinal reflexes with low doses (4-16 mg in this study). We developed an acute sedation neurobehavioral scale for NHP (Supplemental Figure S2C), where reflexes were assessed as a part of standard neurological exams 3-4 hours post dosing, and scores are derived post-hoc from the reports as described in Materials and Methods. In NHP, animals were dosed weekly with increasing doses of ASO (4, 8, 12 and 16 mg). As in mice and rats, ASO6 had a dose dependent acute sedation response in NHP (top dose mean score 5, p < 0.01, Figure 3B). Overall, increasing doses of the same ASO caused increasing levels of acute sedation in mice, rats, or NHP following either ICV or IT administration.

### Acute sedation is driven by acute dose level, and does not change with repeated administrations

Next, we investigated the effect of repeated ASO administration on the acute sedation phenotype, to determine if there was an increased response with ASO tissue accumulation, or signs of tolerance after repeat dosing. IT dosing in rats with ASO6 showed a consistent 3-hour acute sedation score across all administrations when the dose level was maintained at either 100 µg (acute sedation score = 0) or 300 µg (acute sedation score = 3), with no statistically significant changes in score with repeated administrations (Figure 4A-B). To determine if this was also true in NHP, the species most relevant for human safety assessments, we again utilized ASO6. In the previous study, 4 mg of ASO did not cause an acute sedation response, but a mild loss of spinal reflexes signal was detected in some animals at 8 mg (Figure 3B). To determine the effects of repeat dosing, we treated animals weekly for 6 weeks with 4 mg ASO, for a total of 24 mg. No acute sedation signal was observed in the NHP with repeated administrations of 4 mg (Figure 4C-D). Together, these data confirm there is no sensitization with repeated administrations, and suggest acute sedation is driven by high local drug concentrations following dosing, rather than by accumulation of drug in tissues following repeat administrations.

**Figure 4.**
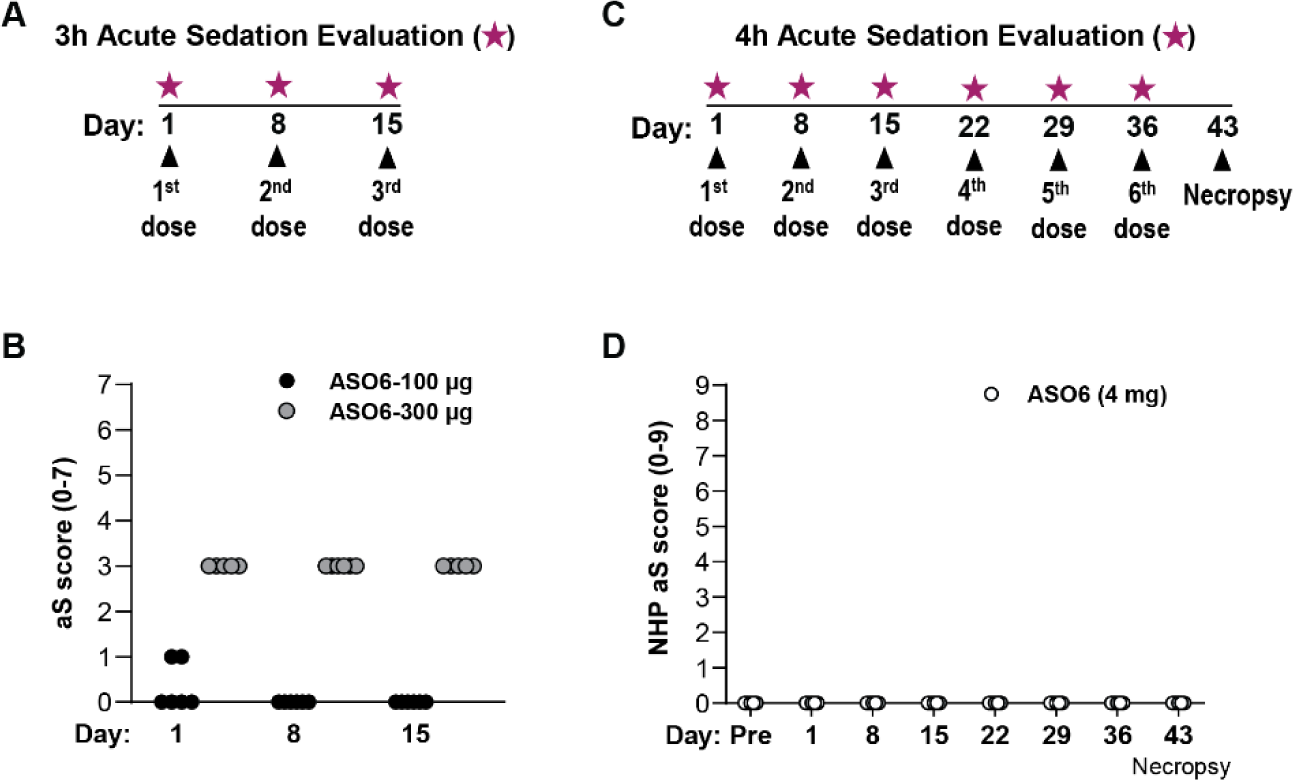
Acute ASO sedation responses do not change with chronic ASO accumulation. (**A**) Experimental design used in (**B**), where rats received 3 doses of ASO (arrowheads) on day 1, 8 and 15 and acute sedation was scored on the scale 3h after each dose (pink stars). (**B**) Rats received either 100 µg (n = 6) or 300 µg (n = 4) of ASO6 via IT administration in rats. The 100 µg dose causes an acute sedation score of 0-1 across all doses and does not increase with successive doses. At 300 µg, ASO6 causes an acute sedation score of 3 across all doses. (**C**) Experimental design used in (**D**), where NHPs were dosed with 4 mg of ASO weekly on day 1, 8, 15, 22, 29, and 36 (arrowheads). NHPs were evaluated at 4h after each dose for acute sedation (pink stars) and necropsy performed a week after the last dose on day 43. (**D**) In NHP, acute sedation phenotypes measured by spinal reflexes are not observed with ASO6 at 4 mg even with repeated administration. Stats for **B** and **D** are nonparametric Kruskal-Wallis test and all time points are non-significant.

### Reducing PS backbone modifications can mitigate acute sedation while maintaining potency, and commonly used 2’ sugar modifications have no effect

Next, we determined if chemical modifications contribute to acute sedation response. PS modifications have been employed in ASO backbones to confer drug-like properties such as increased cellular uptake and metabolic stability [27]. PS modifications are also associated with increased non-specific protein binding [14]. We hypothesized that the PS modification may be contributing to the acute sedation response. In ASOs with sugar modifications in the wings, PS is not needed in the wings as the PO backbone is protected from nucleases by the sugar modifications. To test this hypothesis, we compared acute sedation levels of two versions of ASO 5, a 5-10-5 MOE gapmer targeting mouse Malat1 RNA that is uniformly modified with the PS modification. For ASO5_PO we used the same sequence but did not include the 6 PS modifications in the wings (Fig 5A and 5B; Supplementary Table S1). Both ASO5 and ASO5_PO elicited dose-dependent increases in the acute sedation (Figure 5C), but the magnitude of response was attenuated with ASO5_PO (2 points lower, p > 0.05; Figure 5C). There was no significant difference in the potency of ASO5 and ASO5_PO in reducing Malat1 RNA in the cortex (ASO5 ED_50_ 18.7 µg vs ASO5_PO ED_50_ 27.6, with overlapping 95% CI; Figure 5D), and a small shift in potency in the spinal cord.

**Figure 5.**
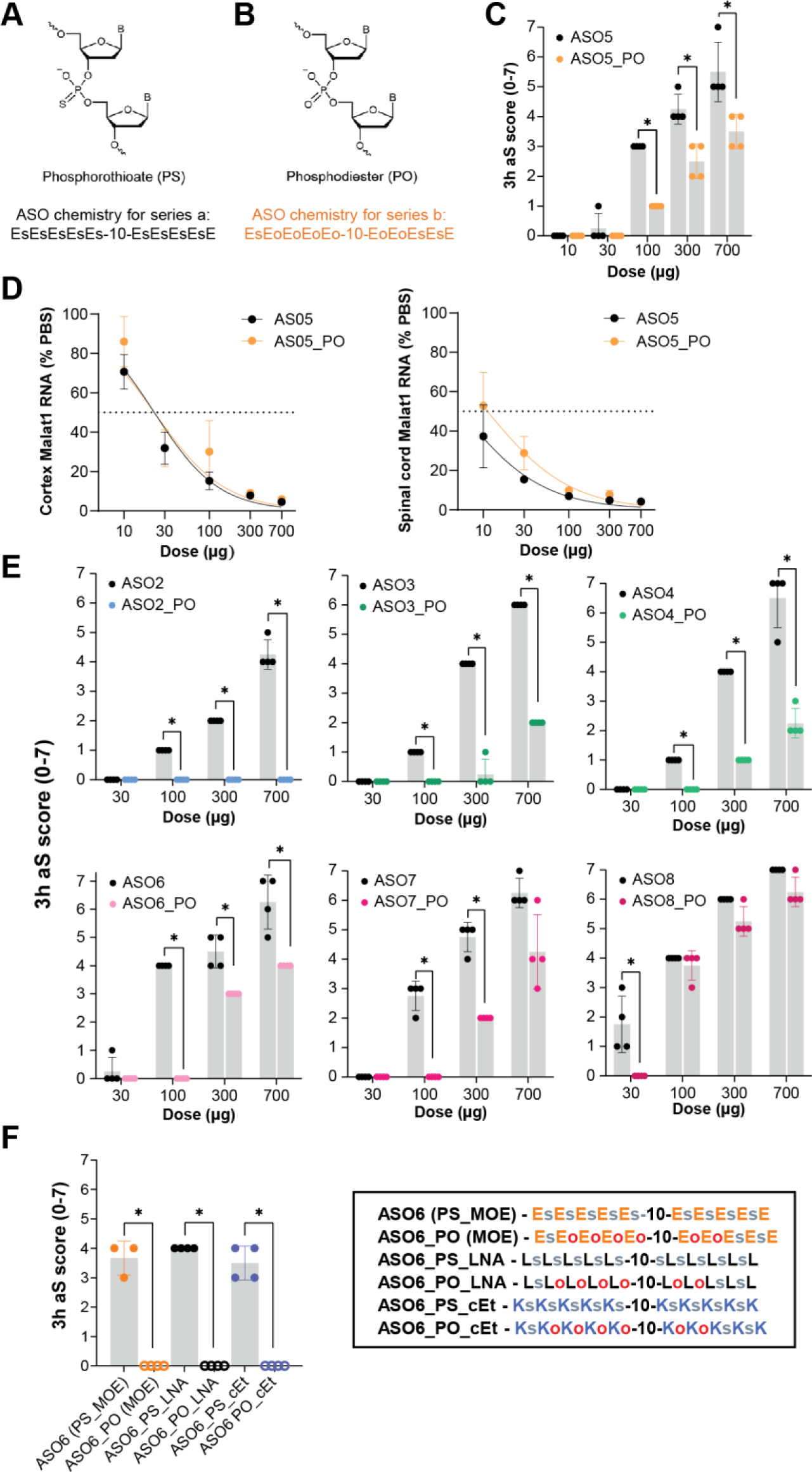
Reducing PS backbone content mitigates acute sedation, but 2’ sugar modifications do not. (**A**) The phosphorothioate (PS) modification replaces a non-binding oxygen on DNA with a sulfur in the phosphate moiety. (**B**) Phosphodiester (PO) is the naturally occurring phosphate moiety in the backbone of DNA. (**C**) ASO5 (fully PS-modified backbone) has significantly higher acute sedation (aS) scores 3 hours post-ICV injection when compared to an ASO of the same sequence containing a mixture of PS and PO in the backbone of the gapmer wing (orange dots). (**D**) Dose response curves 2 weeks post-ICV injection show the potency of a Malat1-targeted ASO with fully modified with PS backbone (ASO5) is minimally affected by modification to a mixed PS/PO backbone (ASO5_PO) with ED50 of 18.7 and 95% confidence interval 15.6 to 22.5 for ASO5 and ED50 of 27.6 and 95% confidence interval 18.7 to 40.7 for ASO5_PO in the cortex. (**E**) 3h acute sedation scores across multiple ASO sequences synthesized with either full PS (black) or mixed PS/PO (colors) backbone show backbone chemistry affects acute sedation severity in most, but not all sequences. (**F**) ASO6 sequences with various sugar ring modifications (MOE, LNA, or cEt) do not affect acute sedation. However, for each of those sugar ring versions, reduced PS linkages mitigated acute sedation entirely. Key: Backbone modifications for PS indicated by **s,** and PO is indicated by **o.** For 2’ sugar modifications, MOE is indicated by **E**, LNA indicated by **L**, cEt indicated by **K.** Stats for **C, E** and **F** are nonparametric Mann-Whitney test and (**D**) is Non-linear Regression (Motulsky). Each data point represents the mean and error bars represent standard deviations. *P < 0.05, **P < 0.01, ***P < 0.001, ****P < 0.0001.

The same strategy was applied across an additional six 5-10-5 MOE gapmer ASO sequences (ASOs 2-8; Table 1), where PS modifications were not included in 6 linkages in the wings (Supplementary Table S1). Reduction of PS in the backbone conferred a range of reduction in acute sedation scores 3 hours after 30-700 µg ICV injection in mice, from complete abolition with ASO2, to highly significant reductions with ASO3 and ASO4, to no significant reduction with ASO8 (Figure 5E). These results indicate reducing PS content can reduce acute sedation in most sequences, but not all.

Gapmer ASOs often incorporate either MOE, LNA, or cEt 2’ sugar modifications to improve binding affinity to target RNA sequences [28]. We generated different versions of ASO6, a 5-10-5 20-mer with either five MOE, LNA, or cEt sugar modifications in the 5’ and 3’ wings and either a full PS or a PO/PS mixed phosphate backbone (Figure 5F; Supplementary Table S1). Mice were injected via ICV with 100 µg ASO and acute sedation was quantified at 3 hours post-injection. Each of the full PS modified versions elicited acute sedation scores of 3-4, regardless of sugar ring chemistry, while scores for each of the PO/PS versions were zero. These data indicate 2’ sugar ring chemistries do not affect acute sedation, but phosphate linkage chemistry can significantly change acute sedation, regardless of sugar ring modifications.

### Acute sedation is not mitigated by divalent cation supplementation in rodents

Previous reports have found divalent cation supplementation in ASO or siRNA dosing solutions can mitigate acute responses, but it is unclear whether activation or sedation phenotypes are affected [10, 15, 18]. Here, we investigated how acute sedation is affected by divalent cations (Ca^2+^ or Mg^2+^) in the dosing solution of ASOs delivered via ICV injection in mice. Animals were treated with aCSF containing 1-100 mM Ca^2+^ or Mg^2+^ without or with an ASO known to cause an acute sedation response (ASO6). Animals were observed from 0-3 hours post-dose, and acute sedation was scored at 3 hours post-dosing. No acute sedation response was observed with the vehicle alone and the acute sedation response was not altered with either Ca^2+^ or Mg^2+^ supplementation (Fig 6A-B), in mice. Of note, all animals treated with 100 mM Ca^2+^ died immediately at dosing and were not included in the 3 hour post-dose evaluation (Figure 6B). Of note, the solution was not controlled for tonicity and may contribute to the effect of high IT Ca^2+^ alone. These results indicate that the acute sedation response in rodents described here is not mitigated by divalent cation supplementation. However, extremely high levels of Ca^2+^ can be lethal when delivered to the CSF and may confound interpretation of acute tolerability studies with ASOs or siRNAs.

**Figure 6.**
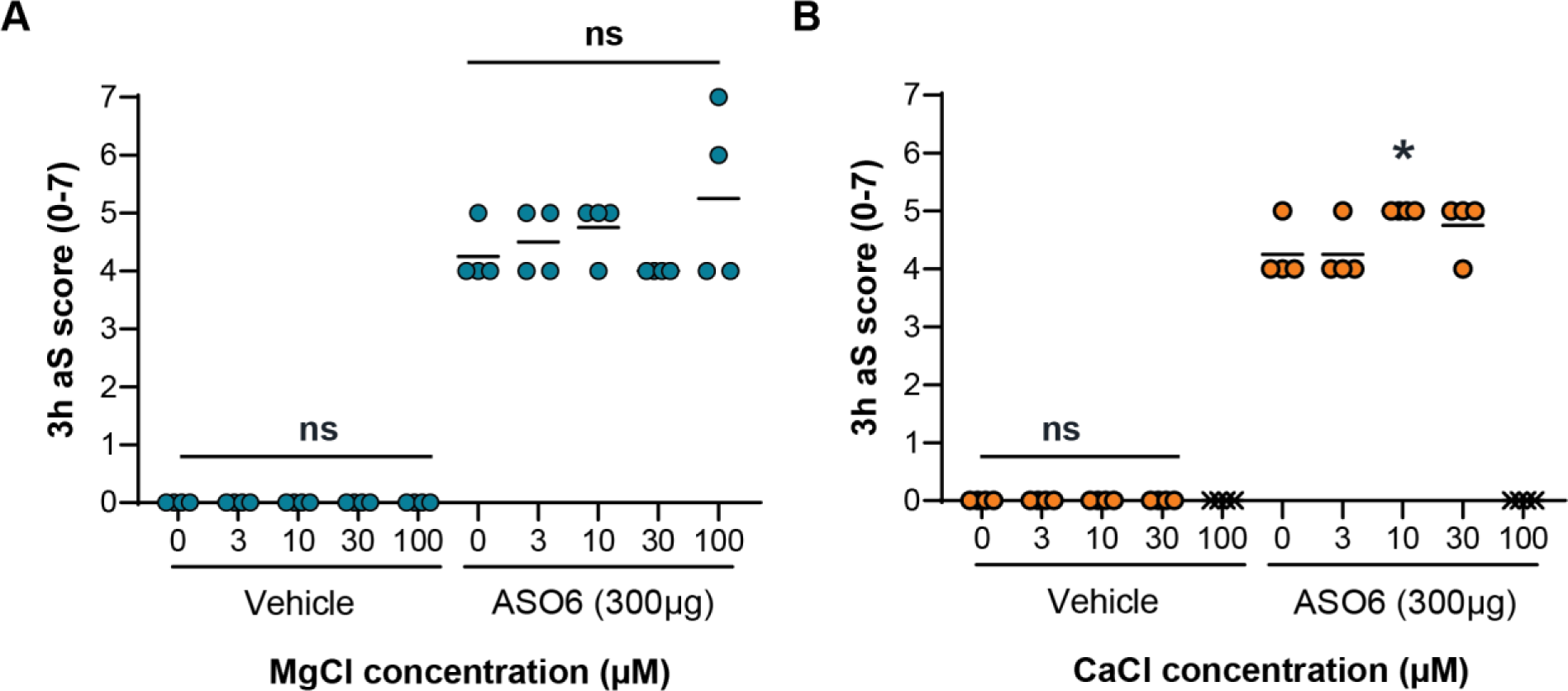
Divalent cation supplementation in ASO dosing solutions does not mitigate acute sedation (aS). (**A-B**) Increasing concentrations (0-100 mM) of (**A**) Mg2+ (blue) or (**B**) Ca2+ (orange) were administered in aCSF via ICV injection in mice either alone (vehicle) or with 300 µg ASO6. Acute sedation level was assessed 3h after injection with no to minimal effect of cation supplementation on the ASO acute sedation response. Each data point represents individual animals’ scores. *P < 0.05, **P < 0.01, ***P < 0.001, ****P < 0.0001 by the nonparametric Kruskal-Wallis test and Dunn’s multiple comparisons post-hoc test. Note: x = died immediately at dosing. Multiple comparisons were done against the 0 mM cation level in each condition (+ or - ASO).

### Acute sedation response is correlated with inhibition of motor circuit neuronal transmission

To determine if acute sedation is caused by inhibition of neuronal transmission, we evaluated activity of neural pathways *in vivo* in rats. Specifically, we recorded activity of motor and sensory pathways at baseline, 2 hours, and 24 hours following IT administration of ASOs in rats. We selected a panel of 5 ASOs that had a range of acute sedation scores after a 700 µg IT dose in rats: 0 (no sedation) for ASO2_PO and ASO3_PO, 1 (tail/foot paresis) for ASO4_PO, 3 (tail and hindlimb paresis) for ASO7_PO, and 4 (tail, hindlimb and trunk paresis) for ASO8_PO. As the acute sedation response is characterized by loss of motor activity, we first evaluated the full motor circuit from the CNS to the peripheral muscle, by quantifying myogenic motor evoked potentials (MMEP) following transcranial electrical stimulation of the motor cortex and recording in the right hind paw muscle (Figure 7A). Consistent with the lack of a behavioral response, no inhibition of the MMEP occurred with non-sedative ASOs 2_PO and 3_PO. In contrast, ASOs 4_PO, 7_PO, and 8_PO caused complete reduction of MMEP 2 hours post-IT injection, indicating full suppression of signaling reaching posterior muscle after cortical stimulation. All MMEP recovered by 24 hours after dosing. This is consistent with the inhibition of distal motor activity at 2 hours for all three ASOs (4_PO, 7_PO, and 8_PO) and restoration of motor behavior at 24 hours.

**Figure 7.**
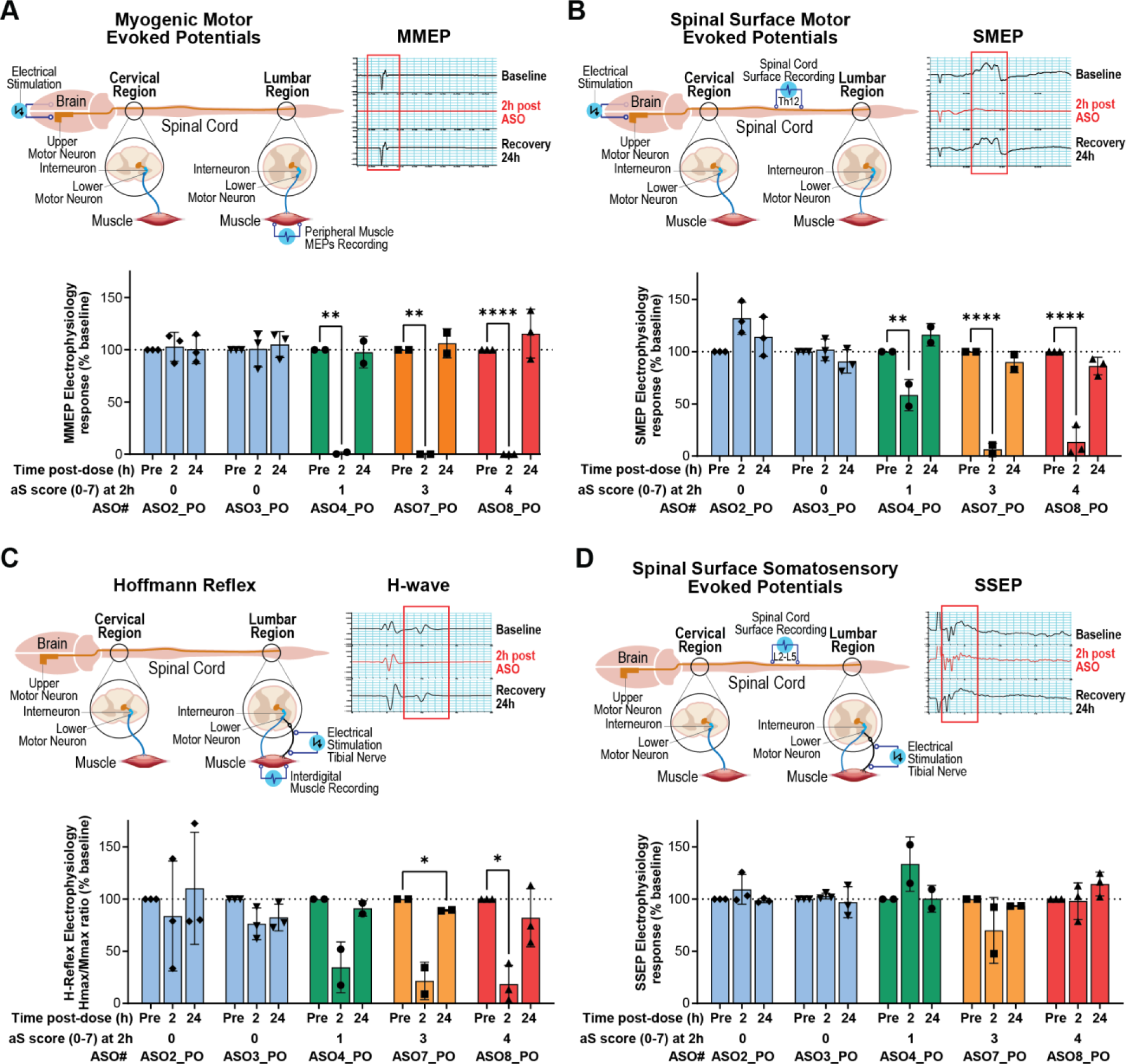
*In vivo* electrophysiology following IT administration of some ASO sequences shows transient motor pathway inhibition in the CNS. Rats were dosed intrathecally with vehicle (PBS) or 700 µg ASO. Electrophysiological recordings were taken at baseline, 2h post-ASO dose, and 24h post-ASO dose. The blue bars represent non-acutely sedative ASOs, the green bars represent a mildly sedative ASO, and the orange and red bars represent moderately acutely sedative ASOs. Upper left panels of A-D are schematics of each experimental set up. Upper right panels of A-D are representative traces from the most severely sedative ASO8_PO at 2h post-dose. (**A**) For myogenic motor evoked potentials, stimulation occurred in the motor cortex and motor pathway activity recordings in the hind paw muscle. Non-sedative ASOs did not affect activity. All acutely sedative ASOs (4-8) abolished activity. (**B**) For spinal surface motor evoked potentials, stimulation occurred in the motor cortex and output was recorded in the thoracic (T12) spinal cord corticospinal tract. Acutely sedative ASOs suppressed activity in a severity-dependent manner at 2h post-injection with restoration of firing by 24h. (**C**) For Hoffman reflex recordings, stimulation occurred in the tibial nerve and recording in the hind paw muscle. Only moderately acutely sedative ASOs affected the monosynaptic spinal reflex. (**D**) For spinal surface somatosensory evoked potentials, stimulation occurred in the tibial nerve and recording in the lumbar spinal cord. No ASO affected somatosensory activity. Bars represent the mean and error bars represent standard deviations. *P < 0.05, **P < 0.01, ***P < 0.001, ****P < 0.0001 using two-way ANOVA and Dunnett’s post hoc test.

Behaviorally, the more severe the acute sedation response, the further anterior the animal is affected following IT administration. To assess a more anterior spinal cord response, transcranial electrical stimulation of the motor cortex was again used, but this time spinal surface motor evoked potentials (SSMEP) were measured on the spinal surface over the exposed corticospinal tract at thoracic level 12 (Figure 7B). Again, non-sedative ASOs 2_PO and 3_PO had no effect on the SSMEP. The mildly acutely sedative ASO4_PO elicited a partial reduction, and moderately sedative ASOs 7_PO and 8_PO caused a nearly complete reduction in SSMEP 2 hours post-IT administration, with recovery by 24 hours. Consistent with the behavioral response, the effect at this cord level was sequence specific. For each ASO, the severity of SSMEP suppression was reflected in the severity of the acute sedation score.

We also measured the Hoffman reflex (HR; a proprioceptive reflex) to evaluate the local lumbar motor circuit in more detail. The tibial nerve was stimulated, and we recorded in the right hind paw muscle (Figure 7C). The tibial nerve projects to the α-motor neuron in the spinal cord (synapse from sensory neuron to motor neuron), which projects to the muscle in the periphery, where the HR is recorded. ASO2_PO and 3_PO had no effect on the HR. ASO4_PO caused a non-statistically significant reduction in HR and ASOs 7_PO and 8_PO caused significant reduction of the HR 2 hours post IT administration. These results reflected the severity of the acute sedation score of each ASO. The HR was fully recovered by 24 hours in all cases. These results demonstrate that acutely sedative ASOs interrupt a monosynaptic reflex involving stimulation of sensory fibers, a synapse in the spinal cord, and response in α -motor neurons.

Finally, to ask whether sensory fibers themselves were impacted by acute sedation, somatosensory evoked potentials were measured by stimulating the tibial nerve, but now recording on the lumbar spinal cord surface (L2-L5) to evaluate sensory afferents (Figure 7D). No ASO had any effect at 2 or 24 hours after intrathecal dosing on sensory afferent response, indicating intact sensory signaling from the periphery into the central nervous system. This is consistent with observations during acute sedation that when the toes of the animals are pinched, the animals react by abdominal twitching or turning their head towards the stimulus, even if more posterior regions such as the foot itself may not be capable of moving at the time.

Taken together, the acute sedation response is likely caused by inhibition of local spinal cord synapses, likely due to high concentrations of ASOs in the hours post-intrathecal dosing.

### Acute sedation phenotypes resolve with clearance of ASO from bulk CNS tissue and the extracellular space

To examine the kinetics of acute sedation as it relates to ASO intracellular uptake and clearance, rats were administered ASO6 (full PS backbone) or ASO6_PO (mixed PS/PO backbone) and acute sedation scores determined prior to necropsy at 1, 3, 6, and 24 hours post-IT administration. ASO tissue concentration was quantified, and ASO localization was evaluated by immunohistochemical staining for ASO. Consistent with previous results (Figure 5C, 5E-F) acute sedation scores were higher with the full-PS backbone version ASO6 than with ASO6_PO, peak sedation scores occurred between 3-6 hours, and resolution of acute sedation by 24 hours post-dose (Figure 8A). ASO concentrations in bulk tissue were similar for ASO6 and ASO6_PO and peaked at 3 hours post-IT injection during the peak of the acute sedation response (Fig 8B). ASO staining using an anti-ASO antibody followed the same overall pattern of peak distribution into the parenchyma (Figure 8C). In the ventral horn, where alpha-motor neurons reside, at 3 hours post-dose, ASO is present both inside and outside of neurons (Figure 8D). By 24 hours ASO has cleared from the extracellular space, and much of it from the CNS, with the ASO that is retained in the CNS mostly localized intracellularly (Figure 8D insets).

**Figure 8.**
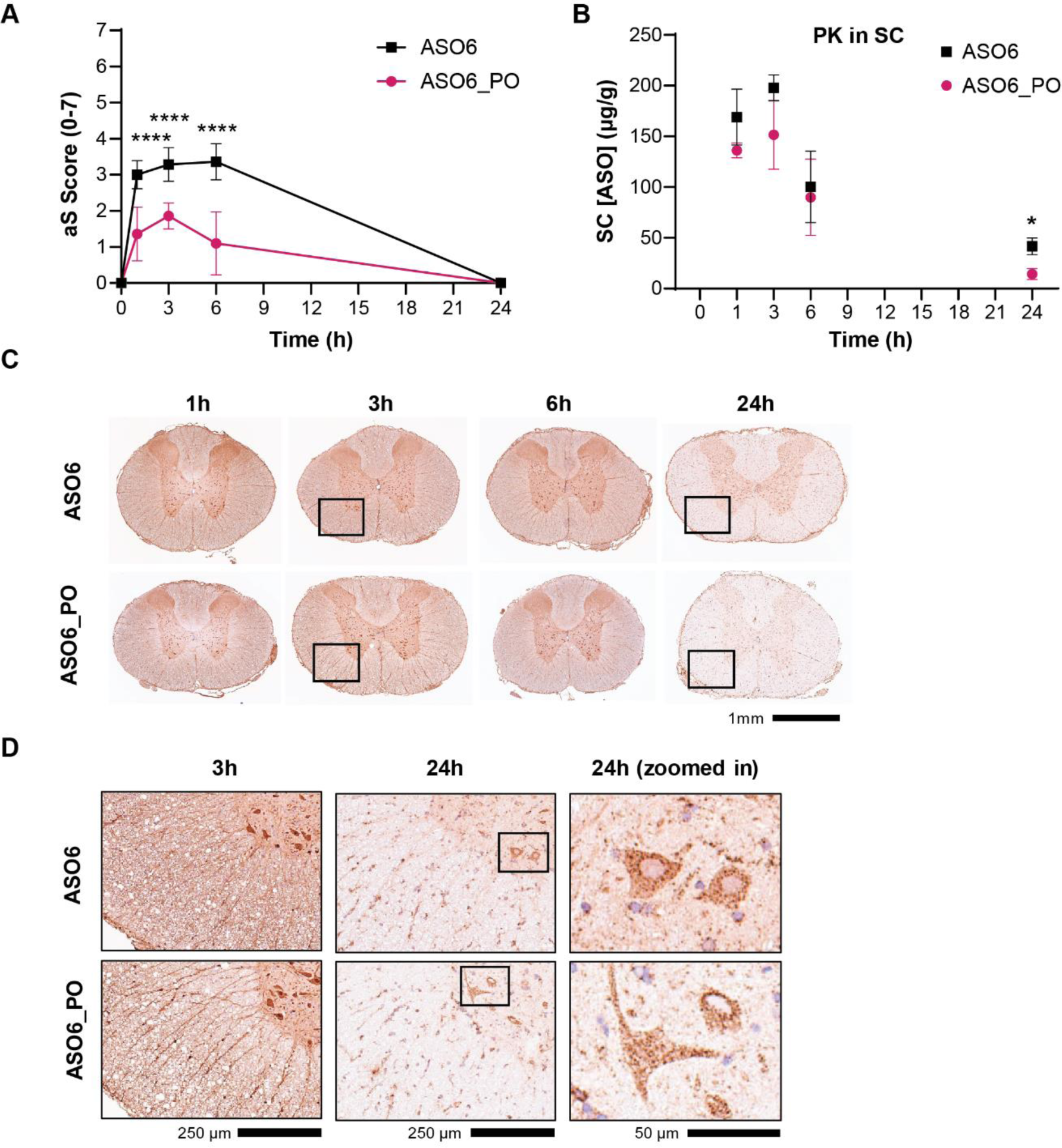
Acute sedation responses resolve with ASO clearance and cellular internalization. (**A**) Time course of acute sedation (aS) responses in rats injected intrathecally with either a fully-PS modified version (ASO6) or a mixed PS/6PO backbone version (ASO6_PO; Table S1) of the same ASO sequence. ASO6_PO acute sedation peaked at a lower score and began to resolve sooner than ASO6 with the response to both versions fully resolving by 24 hours. (**B**) ASO concentration over time in the spinal cord (SC). (**C-D**) ASO staining in the ventral horn of the spinal cord shows strong extracellular distribution of both ASOs during peak acute sedation (3 hours post-injection, left panels). By 24 hours, most ASO has been cleared from extracellular space or internalized (middle panels; inset of staining in motor neurons in right panels). Each data point represents the mean, error bars represent standard deviations, and *P < 0.05, **P < 0.01, ***P < 0.001, ****P < 0.0001. Stats for **A** and **B** using nonparametric Mann-Whitney test.

### Acutely sedative ASOs suppress spontaneous activity in primary cortical cultures

To further investigate underlying mechanisms of acute sedation, we established an *in vitro* model using MEA recordings of spontaneous neuronal activity in primary mouse cortical cultures. MEA assays are routinely used to explore neurotoxicity of drug compounds [29]. Following isolation, cultures were allowed to mature for 2 weeks to form neuronal networks that produced measurable spike rates, bursting patterns, and network bursts on the MEA assay (data not shown). As the sedative response to ASOs is dose- and sequence-dependent *in vivo* we set out to first determine if this was replicated *in vitro*. We selected 3 ASOs, one with zero, one with moderate, and one with severe acute sedation response *in vivo* following a 300 µg ICV dose in mice. The ASOs inhibited network firing relative to baseline in a dose- and severity-dependent manner (Figure 9A). We used this information to select a single *in vivo* dose level (300 µg) and an *in vitro* concentration level (50 µM) to determine if there is a correlation between *in vivo* and *in vitro* response, independent of sequence of chemistry. To test this, a panel of 33 ASOs was evaluated with varying lengths and chemistries, including MOE, LNA and cEt sugar modifications that are often used ASO designs [30, 31]. The *in vivo* acute sedation score was highly correlated with the level of firing inhibition (r = 0.84; Figure 9B), indicating the *in vitro* assay can capture the degree of acute sedation of a particular sequence, regardless of chemistry.

**Figure 9.**
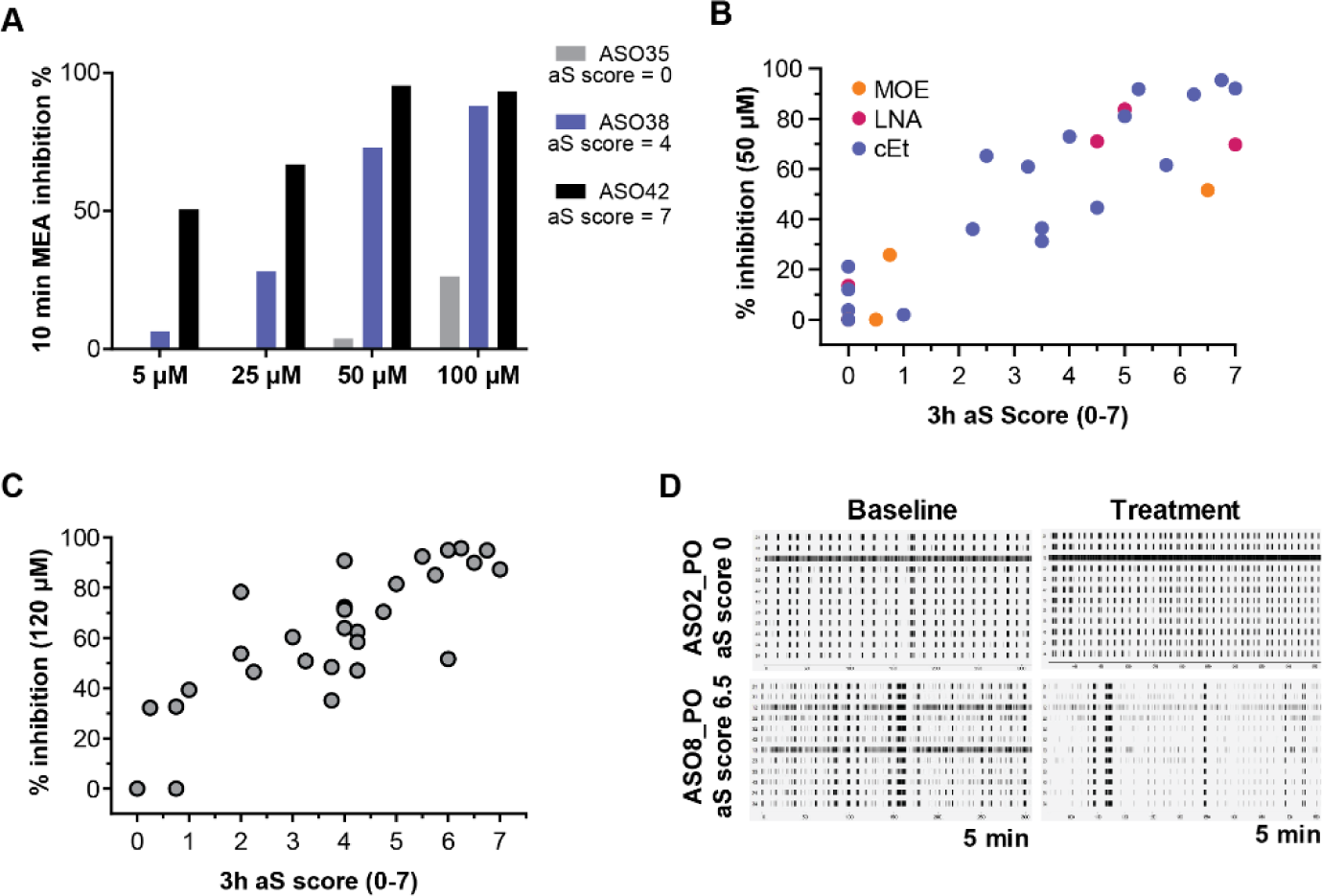
*In vivo* ASO-mediated sedation scores correlate with inhibition of neuronal firing *in vitro*. (**A**) Dose response studies with primary neurons treated ASO35, ASO38, and ASO39 at 5 µM, 25 µM, 50 µM, and 100 µM dose of ASO showed that 50 µM ASO concentration reflected the acute sedation (aS) scores found after a 300 µg ICV injection in mice (ASO35 = 0; ASO38 = 4; ASO39 = 7). Lower concentrations did not show maximal inhibition of firing with the most severe ASO42; higher concentrations did not separate the moderate ASO38 from severe ASO42 (**B**) Percent inhibition of firing in primary neurons treated with 33 different ASOs (7 LNAs, 21 cEts and 5 MOE gapmers) at a concentration of 50 µM is highly correlated to 3h acute sedation scores after a 300 µg ICV injection in mice (r = 0.84 with 95% CI = 0.69 to 0.92), regardless of chemical modification and sequence length (Supplementary Table S2). (**C**) Primary neurons treated with MOE gapmer ASOs (n=34) at 120 µM concentration showed a strong correlation (r = 0.9 with 95% CI = 0.79 to 0.95) between % inhibition of firing rate and 3h acute sedation score following a 700 µg ICV injection in mice. (**D**) Representative raster plots of neuronal spike patterns at baseline and after treatment with 120 µM of an ASO that has a high acute sedation score of 6.5 shows visual suppression of the *in vitro* firing rate. Stats for (**B**) and (**C**): nonparametric Spearman r correlation test.

Given the dose dependence of the response both *in vitro* and *in vivo*, we hypothesized the assay could be tailored to assess ASOs at different dose levels to establish the desired therapeutic index. To identify ASOs with the lowest potential for acute sedation (as described in Figure 6), we selected a panel of ASOs with lower PS content and we increased the dose at which we evaluated acute sedation *in vivo* to 700 µg. Here we were able to identify ASOs that still had little acute sedation at 700 µg in mice. We also correspondingly increased the concentration at which we tested the ASOs *in vitro* to 120 µM after a concentration-ranging study (Supplementary Figure S3) similar to our previous exercise with a new set of mild, moderate, and severely sedative ASOs (Figure 9A). Again, we saw a strong correlation between the *in vitro* and *in vivo* response (r = 0.9; Figure 9C). Screening ASOs *in vitro* at 120 µM and selecting compounds with < 50% inhibition of baseline firing rate should identify ASOs with little to no potential for acute sedation at 700 µg in mice, the human equivalent dose based on CSF scaling of 2450 mg (based on reported CSF volume ranges [32]).

### Suppression of spontaneous firing is reversed by removal of ASO from culture media

The kinetics of the acute sedation response and ASO distribution *in vivo* suggest that ASO in the extracellular space is potentially responsible for the inhibition of neuronal firing. To test this directly, we asked if ASO-mediated inhibition of spontaneous firing *in vitro* would be reversed by removing the ASO from the media, and thus the extracellular space. To do this we applied either media, or a panel of ASOs with different levels of *in vivo* acute sedation to the primary cultures and recorded for 5 minutes at baseline, during ASO treatment, and after washout of ASO from the culture media (Figure 10A-D). As before, ASOs with zero *in vivo* acute sedation had no effect on the *in vitro* firing rate while ASOs with acute sedation fully suppressed firing during treatment, and those with moderate acute sedation moderately inhibited *in vitro* firing rates (Figure 10A-D). Following washout, firing rates were normal for all ASOs and no differences were observed amongst treatment groups. These results indicate that the inhibitory effect of ASO fully reversed with removal of ASO from the extracellular space.

**Figure 10.**
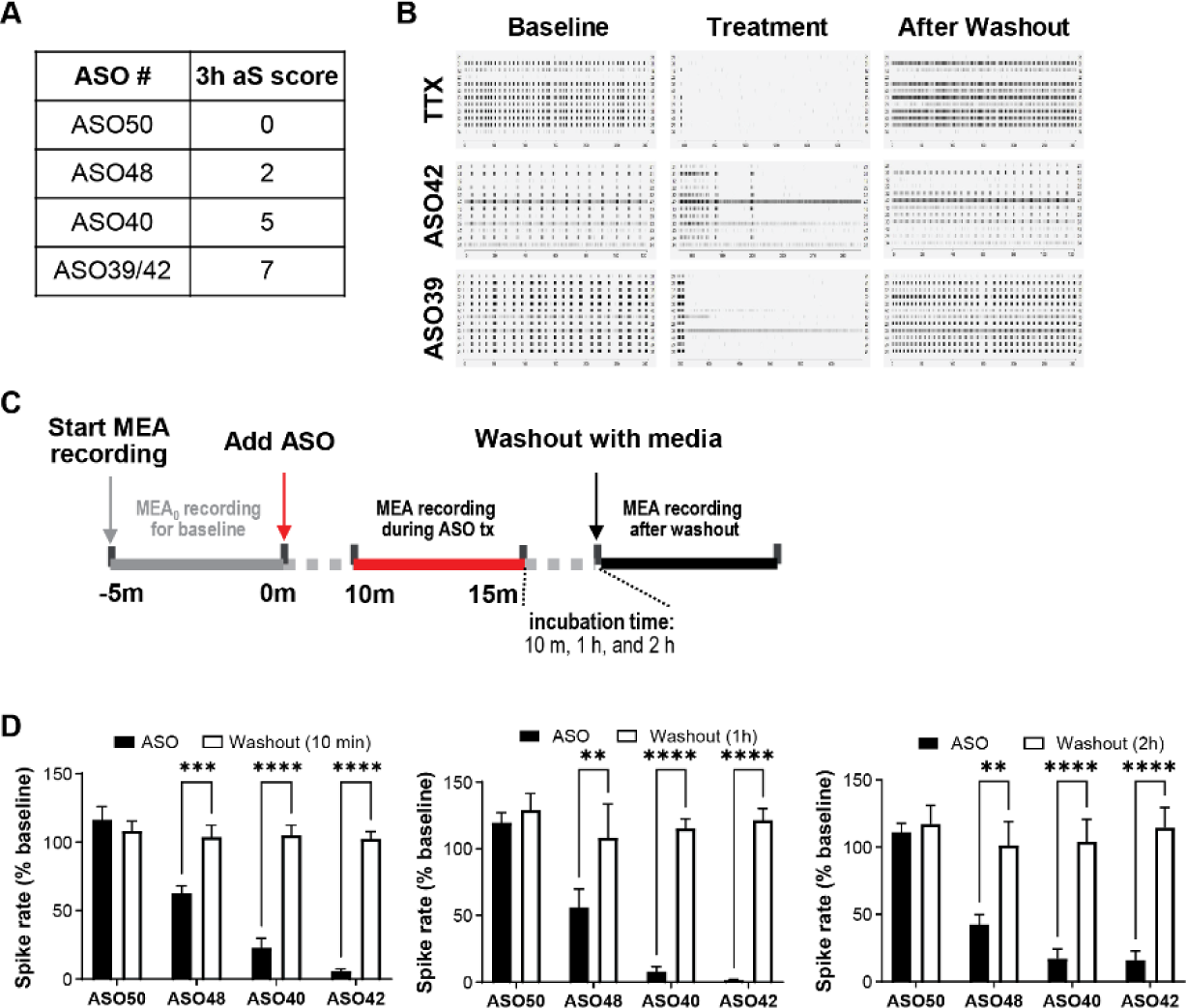
ASO-mediated inhibition of firing in primary cultures is reversible upon washout. (**A**) Table of *in vivo* acute sedation scores for ASOs used in (**D**) with the corresponding *in vivo* acute sedation score following 700 µg ICV injection in mice. (**B**) Representative raster plots showing primary neural culture firing at baseline, after treatment with TTX (positive control of neurotransmission inhibition), an ASO that causes an *in vivo* 3h sedation score of 6.75 (ASO42), and an ASO that causes an *in vivo* acute sedation score of 7 (ASO42), and after 2 washes with media to remove the treatment. (**C**) Schematic of experimental design with baseline recording for 5 min, addition of ASO with acclimation period of 10 min followed by recording for 5 min, then incubation with ASO for 10 min, 1h, and 2h before washing and recording (5-10 min). (**D**) MEA firing rates after treatment with ASO50, ASO48, ASO40, ASO42 (black bars) and following washout of the ASOs at 10 min, 1h, and 2h post-treatment (white bars) indicates the suppression of firing is reversible immediately after removal of ASO from the extracellular space, regardless of incubation time allowed. Each data point represents the mean, error bars represent standard deviations, and *P < 0.05, **P < 0.01, ***P < 0.001, ****P < 0.0001. Stats for **D**: two-way ANOVA and Šídák’s post hoc test.

### ASO-mediated suppression of neuronal firing can be overcome by any excitatory stimulus

Neuronal transmission can be inhibited by blocking axon propagation, preventing neurotransmitter release into the synapse, or preventing receipt of that signal on the post-synaptic density. The *in vivo* recordings suggest neuronal inhibition is occurring at the synapse and likely does not involve axonal propagation. To further exclude axonal propagation, and to determine if a single receptor or pathway is involved, we applied various neurotransmitters, excitatory agonists, or inhibitory antagonists in dose response during ASO treatment in primary neural cultures. First, in a positive control experiment, tetrodotoxin (TTX) was used to inhibit axonal propagation via blockade of voltage-gated Na^+^ channels. About 90% inhibition of firing was observed, and no neurotransmitter, agonist, or antagonist was able to rescue the firing (Supplemental Figure SF4). This is what is expected if axonal propagation is inhibited. In contrast, ASO8_PO (acute sedation score = 7 at 700 µg via ICV in mice) inhibited firing by 85-90%, which was fully rescued with 100 µM glutamate or 50 µM acetylcholine and partially rescued by NMDA, AMPA, or bicuculine (Fig 11A-E). ASO7_PO (acute sedation score = 4 at 700 µg via ICV in mice) inhibited firing by 60%, which was fully rescued by 50 µM acetylcholine, 50 µM glutamate, 25 µM NMDA, 1 µM AMPA, or 25 µM bicuculine (Figure 11A-E). ASO2_PO (acute sedation score = 0 at 700 µg via ICV in mice) did not result in firing inhibition (Figure 11A-E). As expected, in non-ASO conditions, application of these agents dose-responsively increased firing before inhibiting activity at very high doses (Right panel, Figure 11A-E). These results show ASO-mediated inhibition of neuronal activity could be partially or fully rescued via modulation of any excitatory or inhibitory cues at receptors present in the post-synaptic density. This suggests that ASO likely blocks firing by non-specific protein binding in the synapse. It also suggests ASOs do not inhibit action potential propagation.

**Figure 11.**
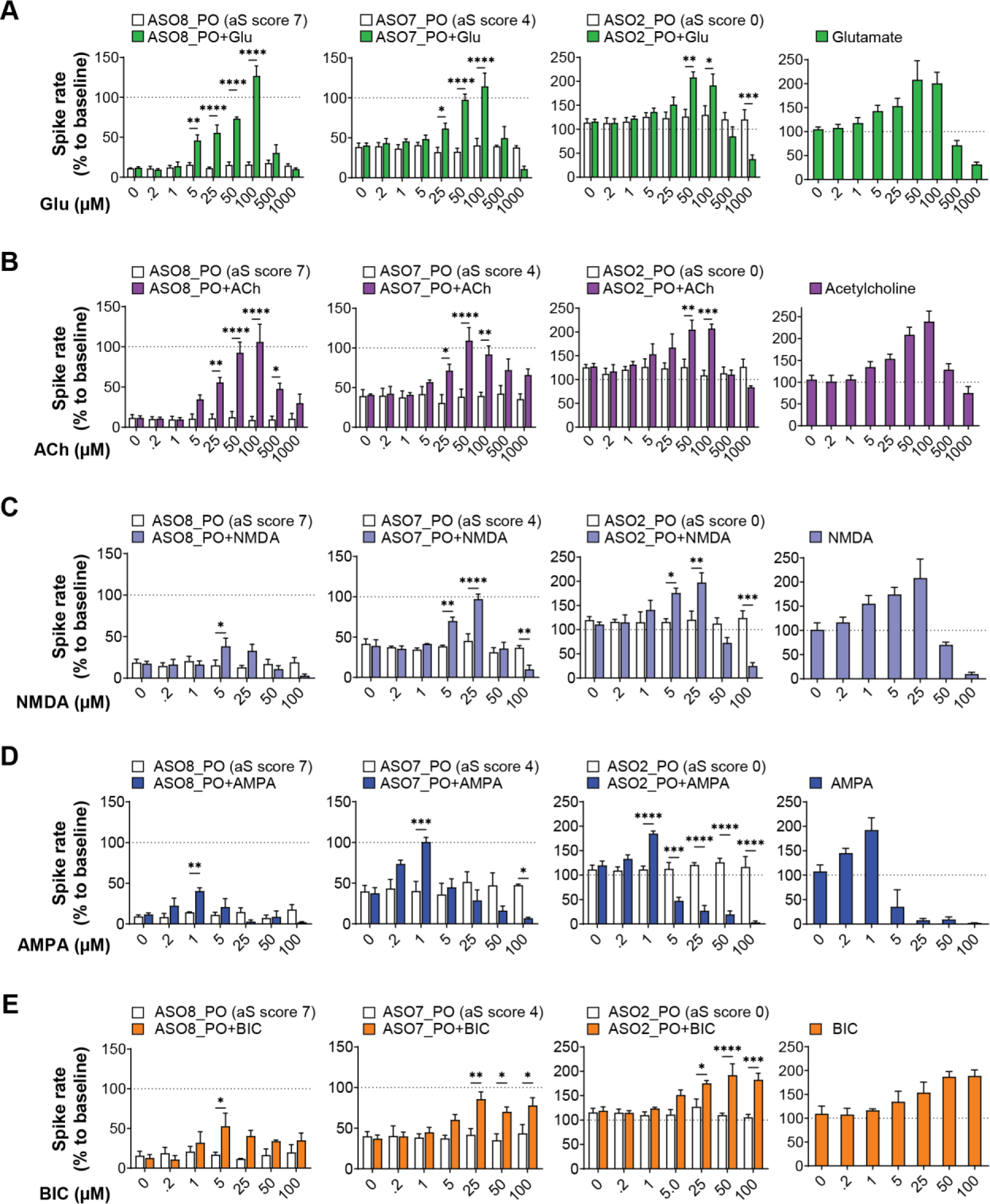
Excitatory agonists and inhibitory antagonists partially or fully rescue ASO-mediated acute inhibition of neuronal firing. (**A-E**) Primary cultures were treated with various agonists or antagonists in a dose response alone (far right panels) or in combination with ASO8_PO, ASO7_PO, and ASO2_PO that cause an acute sedation score of 7, 4, and 0, respectively. Accordingly, ASO8_PO, ASO7_PO, and ASO2_PO caused severe, moderate, and no inhibition in *in vitro* spike rates, respectively. (**A**) Glutamate (Glu) or (**B**) Acetylcholine (Ach) concentrations between 5-100 µM fully rescued any ASO-mediated inhibition of firing. (**C**) N-methyl-D-aspartate (NMDA), (**D**) α-amino-3-hydroxy-5-methyl-4-isoxazolepropionic acid (AMPA), and (**E**) Bicuculline methiodide (BIC) fully rescued moderate inhibition of firing caused by ASO7_PO, but only partially rescued the severe suppression caused by ASO8_PO. Each data point represents the mean, error bars represent standard deviations, and *P < 0.05, **P < 0.01, ***P < 0.001, ****P < 0.0001. Stats for **A**-**E**: two-way ANOVA and Šídák’s post hoc test.

## DISCUSSION

Acute and transient neurobehavioral abnormalities caused by ASOs administered into the CNS have been widely reported [10, 15-18]. Here, we specifically characterize a class of behaviors that involves inhibition of CNS activity, including loss of lower spinal reflexes, hypoactivity, lethargy, paresis, sedation, and ataxia. Following IT administration in rats, acute sedation presents as a loss of corticospinal tract activity and motor function that affects progressively more anterior regions of the body with increasing dose levels or with more severely sedative sequences. Following ICV injection in rodents, we find acute sedation can include loss of locomotor activity and consciousness. In NHP, we observed acute sedation responses including transient loss of spinal reflexes. These behaviors can be captured in acute sedation scales that can be easily adapted to the drug discovery pipeline. The acute sedation response peaks around 3 hours post-administration and is typically fully reversible by 24-hours post-dose, with no identifiable sequelae and no histological correlates.

It should be noted in addition to loss of spinal reflexes in NHP, a wider range of transient, acute sedation responses presenting in the hours after IT ASO administration has been observed [17]. These can include signs such as knuckling (deficits in proprioception positioning manifesting as walking on the digits or knuckles of the hind limbs), ataxia (inability of the truncal, pelvic, and limb muscles to move in unison, creating an unsteady gait), and limb paresis (loss of weight-bearing ability or movement of any limb), and are assessed as a part of standard neurological exams performed post-IT administration of ASOs. We hypothesize the doses used in the NHP studies reported herein were too low to elicit these types of responses.

Our data support a model where the acute sedation response is caused by inhibition of synaptic transmission with high concentrations of extracellular phosphorothioate ASO. The transient nature of the response is coincident with the clearance of ASO out of the extracellular space and into uptake into cells. Acute sedation is a non-specific effect that is highly sequence dependent and can be mitigated by decreasing phosphorothioate content and avoiding G-rich sequences.

A previously unresolved question was how to define the acute sedation response and how the response translates across species, delivery routes, and model systems. We demonstrate the response translates from *in vitro* cell culture to rodent, and from rodent to NHP. It presents differently when the ASO is delivered through either IT or ICV routes to the CSF due to differential local CNS tissue concentrations of ASO. The acute sedation response is distinct mechanistically from the acute activation response [33]. Others who have described acute activation phenotypes such as seizures and hyperactivity have observed these phenomena within the first minutes to one hour following administration with resolution between 2-4 hours [10, 15-18], highlighting an important temporal difference in the evolution of the different acute phenotype classes. Sometimes in these reports, animals are evaluated for acute phenotypes within the first hour but are not reportedly monitored during the peak acute sedation phase (3 hours post-dose). In our experience, there are ASOs that elicit both acute activation and sedation phenotypes, those that elicit one or the other, and those that elicit neither. In the pursuit of discovering safe oligonucleotides for clinical use it is important to evaluate both classes of phenotype immediately after dosing, at 3 hours after dosing, as well as at 24 hours post-dose to ensure an evaluation of all potential acute phenotypes as well as reversibility.

Since most ASOs selected for these studies were not designed to target RNA within the species in which they were tested (mouse, rat, or NHP), on-target cleavage of RNA is not likely to contribute to the neurobehavioral phenotypes observed. Furthermore, the timing of acute sedation within the initial distribution phase of the ASO in first few hours after injection indicates hybridization events are unlikely to contribute. Others have ruled out innate immune activation as causative for any acute phenotype [18].

Our work extends earlier efforts to understand chemical contributions and establishes the severity of acute sedation is dependent on the sequence and backbone chemistry of the ASO, but not the 2’ sugar modifications (either MOE, LNA or cEt). ASOs with identical chemical motifs but different sequences exhibited a wide range of acute sedation levels. We found *in vivo* acute sedation severity was positively associated with enrichment for guanine (G) nucleotides. A previous report also reported a positive association with G-rich sequences in an in vitro assay of ASO-mediated suppression of synchronous neuronal activity [16]. However, though present, we did not find a strong negative association between *in vivo* acute sedation and adenosine (A)-rich sequences, as had been reported in the cell assay. We also found replacement of PS linkages with PO in the wings of a gapmer ASO did not profoundly affect the ED_50_ of target RNA knockdown, but did mitigate acute sedation in most sequences, indicating this can be an approach to improving tolerability of ASOs without affecting efficacy.

We modeled acute sedation in primary cortical cultures by measuring spontaneous activity via MEA. The severity of activity suppression in culture was highly correlated to the severity of the *in vivo* acute sedation scores. The fast onset of suppressed neuronal firing in the minutes after ASO application and reversal in the minutes after washout *in vitro* suggests acute sedation is likely mediated at the extracellular level rather than by mechanisms involving cellular uptake and intracellular trafficking. Others have also demonstrated reduced activity in a spontaneous calcium oscillation *in vitro* assay following application of ASO [16] and reduced intracellular calcium levels in rat primary neurons [15]. These results are consistent with the general suppression of neuronal firing that we have observed, and likely secondary to the general decrease in neuronal activity, and not a direct mechanism on intracellular calcium dynamics. It is also unclear whether the ASOs used in the published assays were acutely sedative or activating (or both). Our data set expands upon the previous work with a significantly larger set of oligonucleotides and chemistries and a range of sequences.

We also demonstrate that the inhibitor effect can be partially or fully overcome by adding any number of excitatory cues (glutamate, acetylcholine, etc.), indicating it is likely not mediated by any one specific class of receptor. It is possible that acutely sedative ASOs instead bind multiple postsynaptic targets in the post-synaptic density or prevent presynaptic release of neurotransmitters. Phosphorothioate ASOs are well-known to bind proteins in a non-specific manner and is consistent with the correlation with the phosphorothioate content [34]. We have also confirmed that the acute sedation response does not correlate with CSF neurotransmitter or glucose levels and is not mediated by inflammation (data not shown). Future research is needed to understand the exact mechanism and synaptic functions involved.

## Supporting information

ORourke_aS Supplement

## ACKNOWLEDGEMENTS

The authors thank Donna Sipe and the vivarium staff, oligo synthesis group, PCR core group, histology core staff, Wanda Sullivan and Tracy Reigle and the creative services team and Linda Fradkin at Ionis Pharmaceuticals Inc., for their technical support.

## AUTHOR CONTRIBUTIONS

Jacqueline G. O’Rourke conceptualization, data curation, formal analysis, investigation, methodology, project administration, visualization, writing-original draft. Gemma Bachmann data curation, formal analysis, investigation, and methodology. Curt Mazur conceptualization, data curation, formal analysis, investigation, methodology, writing-review & editing. Keming Zhou conceptualization, data curation, formal analysis, investigation, methodology for in vitro experiments. Oleksandr Platoshyn in vivo electrophysiological experimental design and data generation. Mariana Bravo Hernandez writing-review & editing. Stephanie Klein histological data generation and analysis of ASO uptake. Jonathon Nguyen generate data for ASO concentration. Sebastien Burel generated data for several graphs. Christine Hoffmaster contributed to NHP studies and data generation, Tom Zanardi to NHP studies and data generation, Paymaan Jafar-nejad conceptualization, validation, writing-review & editing, Martin Marsala in vivo electrophysiological experimental design and data generation, Scott P. Henry supervision, validation, writing-review & editing, Eric E. Swayze supervision, validation, writing-review & editing, Berit Powers conceptualization, resources, supervision, validation, writing-original draft, Holly B. Kordasiewicz conceptualization, data generation, resources, supervision, validation, writing-original draft.

## SUPPLEMENTARY DATA

Supplementary Data are available online.

