## Supplementary material for "Reversable Acute Sedation Response of Phosphorothioate Antisense Oligonucleotides Following Local Delivery to the Central Nervous System": ORourke_aS Supplement

### Supplemental figure 1

**A** **Acute sedation (aS) scoring following IT administration in rodents**

| 3h aS score | Rodent IT injection acute sedation scale |
|---|---|
| **0** | Bright, alert, responsive |
| **1** | No tone/movement in tail |
| **2** | Weak posterior posture |
| **3** | Hind limbs don't support weight, but can still move |
| **4** | Hind paws don't move – full hindlimb paralysis |
| **5** | Weak anterior posture |
| **6** | Fore paws don't move, but animal is still breathing |
| **7** | Death |
| Note: Rats are observed 3 hours after dosing and a score is assigned. | |

**B** **Acute sedation (aS) scoring following ICV administration in rodents**

| 3h aS score | Rodent ICV injection acute sedation scale |
|---|---|
| **0** | Bright, alert, responsive |
| **1** | Gait slightly impaired |
| **2** | Hunched and/or impaired gait |
| **3** | Forward movement only after lift |
| **4** | Any movement after lift |
| **5** | Lateral recumbency with response to tail pinch |
| **6** | No response to tail pinch but still breathing |
| **7** | Death |
| Note: Mice or rats are observed 3 hours after dosing and a score is assigned. | |

**C** **Acute sedation (aS) scoring following IT administration in NHP**

| NHP aS score | Exam | Observation |
|---|---|---|
| **2** | Knee Jerk Reflex | The quick extension of a hind leg is noted following the tapping of the patellar ligament with a hard object: **absence of response = + 1 pt per side** |
| **2** | Cutaneous Reflex | The reaction to a series of needle pricks along each side of the abdomen is recorded: **absence of response = + 1 pt per side** |
| **2** | Proprioceptive Reflex | Each foot is inverted and the ability to right the foot is noted: **absence of response = + 1 pt per side** |
| **2** | Sensory Foot Reflex | The foot is stimulated with a sharp object and an avoidance response is observed: **absence of response = + 1 pt per side** |
| **1** | Tail Reflex | When the tail is touched with a sharp object, the tail movement to avoid the stimulus is noted: **absence of response = +1 pt** |
| **9** | | **Total score (cumulative)** |

**Supplemental Figure 1.** Acute sedation scales for IT or ICV administration in rodents, or IT administration in NHP. Acute sedation scales for IT (**A**) or ICV (**B**) administration in rodents and IT administration in NHP (**C**).

### Supplemental figure 2

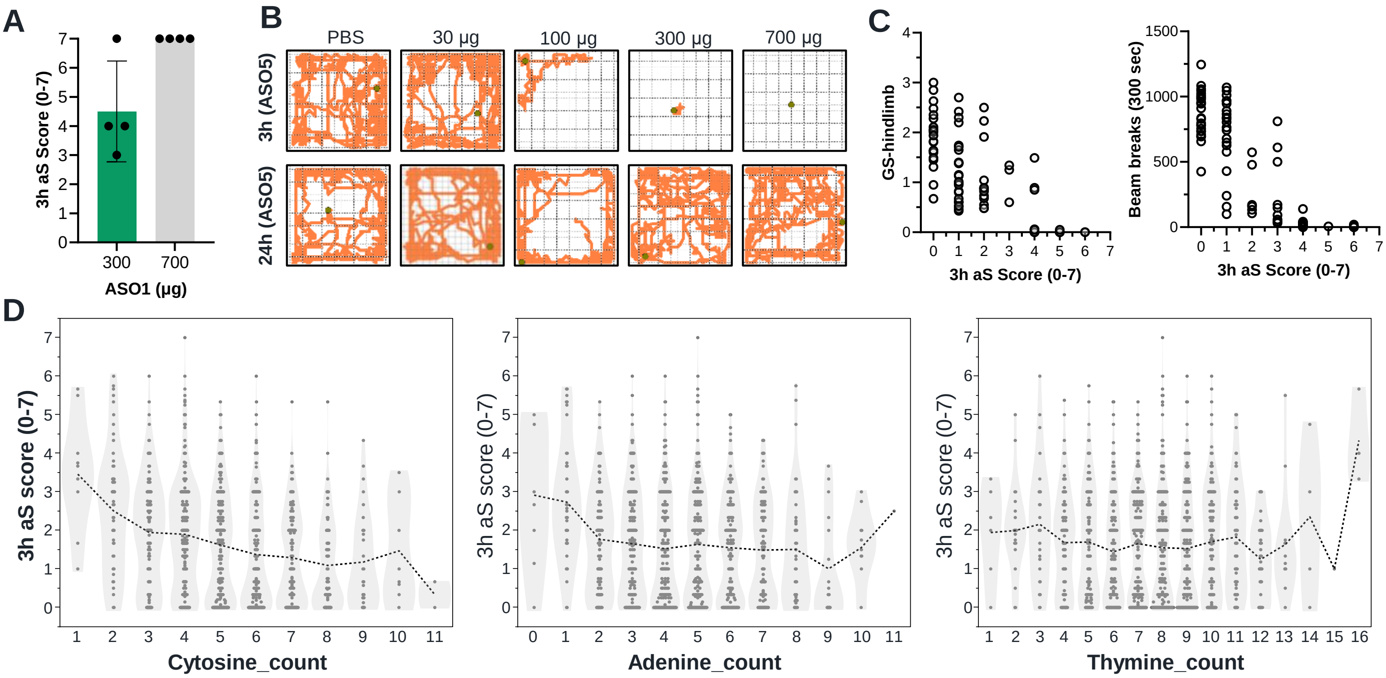


**Supplemental Figure 2.** Acute sedation is sequence dependent and translates to mouse, where measures of grip strength and open field activity correlate with 3h aS scores. (**A**) aS sedation scores 3 hours post-ICV injection of 300 or 700 µg ASO1 in mice as measured on the ICV scale developed for rodents demonstrates the response translates across species from rat (see Fig. 1C) to mouse. Note the 300 µg dose in mice elicited a similar level of acute sedation to the 700 µg dose in rats (Fig. 1C), but 700 µg of ASO1 was lethal for mice. (**B**) Representative OF traces of total activity in mice ICV injected with ASO5 in a dose response study (Fig. 2F) showing dose-dependent suppression of locomotor activity at 3h (upper panels) and recovery by 24h (lower panels) compared to PBS control (far left panels). (**C**) Hindlimb GS is highly correlated between hindlimb GS (r = -0.7611) and OF total activity (r = -0.8463) with acute sedation score. (**D**) Total number of Cytosine (C), Adenine (A), or Thymine (T) nucleotides in a given ASO sequence relative to the 3h acute sedation score *in vivo* following 700 µg ICV injection of ASO in mice. Stats for (C): Nonparametric Spearman r correlation test.

### Supplemental figure 3

A

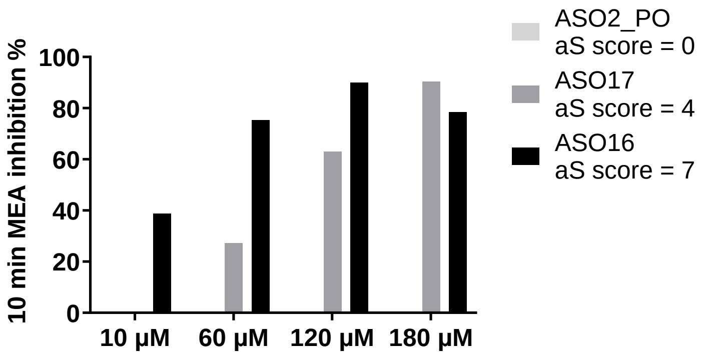


**Supplemental Figure 3.** ASO concentration of 120 µM inhibits neuronal firing *in vitro* to a degree reflecting the severity of *in vivo* acute sedation for different ASO sequences following a 700 µg ASO ICV dose. (**A**) Dose response studies with primary neurons treated with ASO2_PO, ASO17, and ASO8_PO at 10 µM, 60 µM, 120 µM, and 180 µM concentration showed that 120 µM ASO concentration strongly reflected the 3h acute sedation (aS) score after a 700 µg ICV dose in mice, with no inhibition caused by ASO2_PO (3h aS score = 0), 60% inhibition with ASO17 (3h aS score = 4), and 90% inhibition with ASO8_PO (3h aS score = 7). A concentration of 120 µM was therefore selected for follow up experiments.

### Supplemental figure 4

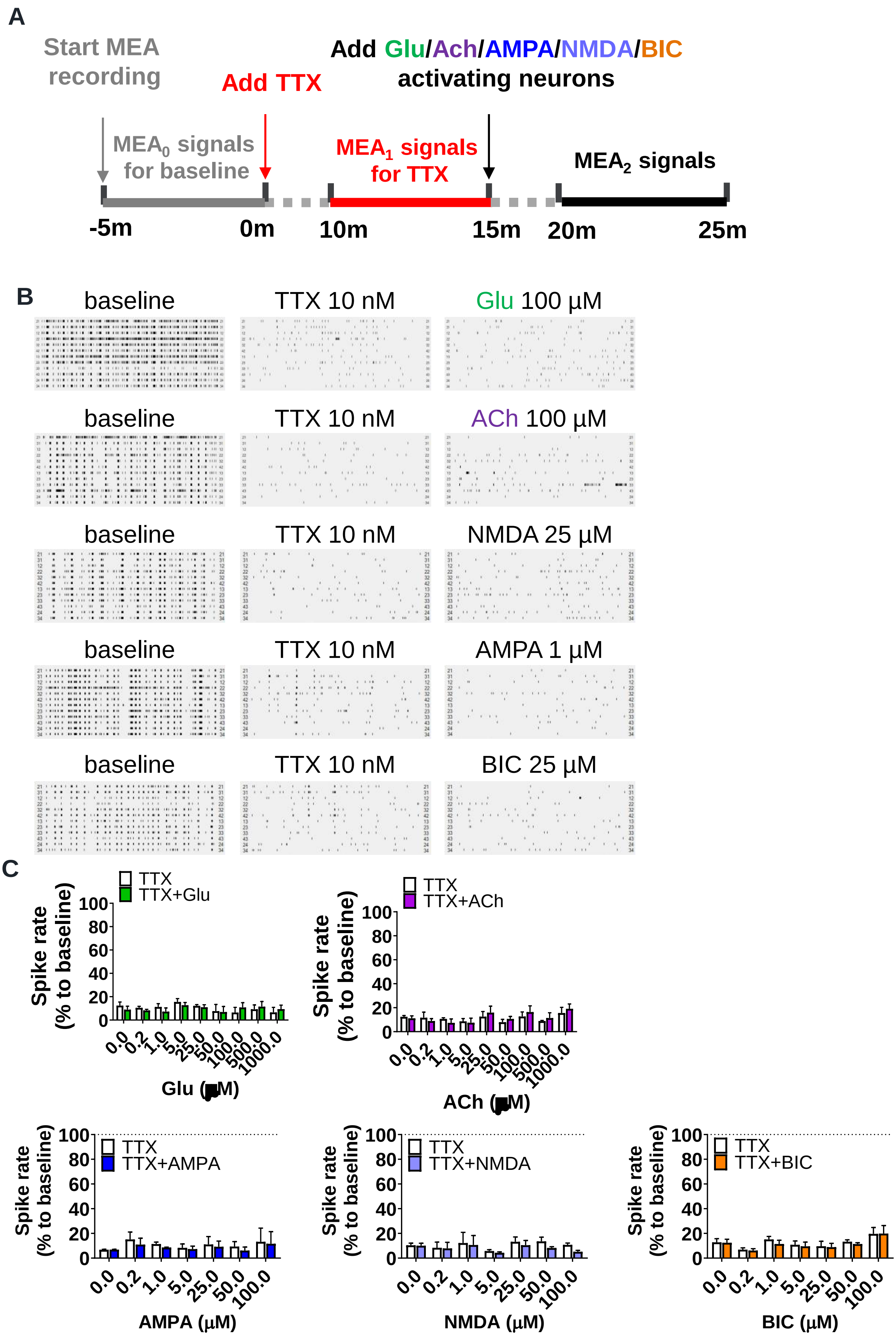

**Supplemental Figure 4.** Application of excitatory neurotransmitters cannot reverse TTX-mediated inhibition of action potential propagation. (**A**) Schematic of the experimental design with baseline recordings for 5 min, 10nM TTX addition and recording for 5 min, then exogenous neurotransmitters added in dose response with recording for an additional 5 min. (**B**) Representative raster plots of neuronal spike patterns at baseline, during treatment with TTX, and after treatment with Glu, Ach, NMDA, AMPA, or BIC. (**C**) Results show ~90% inhibition of neuronal firing upon treatment with TTX, similar to ASO9_PO that causes ~90% decrease in neuronal spike rates compared to baseline control. Neurotransmitter treatment with Glu, Ach, NMDA, AMPA, or BIC could not reverse TTX-mediated inhibition at any concentration.

### Supplemental Table 1

**ASOs used in *in vivo* studies**

| ASO # | SimpleSequence | Figure # |
|---|---|---|
| ASO1 | **TCTCTATTGCACATTCCAAG** | 1 |
| ASO2 | **GTTTTCAAACACACCTTCAT** | 2A-C; 6E; 7A-D |
| ASO2_PO | **GToToToToCAAACACACCToToCAT** | 6E |
| ASO3 | **GTTTTCTTATTTTTAGCATT** | 2A-C; 6E; 7A-D |
| ASO3_PO | **GToToToToCTTATTTTTAGoCoATT** | 6E |
| ASO4 | **CACTTCGAACTCCTGGCGGG** | 2A-C; 6E; 7A-D |
| ASO4_PO | **CAoCoToToCGAACTCCTGGoCoGGG** | 6E |
| ASO5 | **GCCAGGCTGGTTATGACTCA** | 2D-F; 6C-D,F |
| ASO5_PO | **GCoCoAoGoGCTGGTTATGAoCoTCA** | 6E |
| ASO6 (PS MOE) | **CGAGACAGTCGCTTCCACTT** | 6E,F, 8A-D |
| ASO6_PO (MOE) | **CGoAoGoAoCAGTCGCTTCCoAoCTT** | 6E,F; 8A-D |
| ASO6_PS_LNA | **CGAGACAGTCGCTTCCACTT** | 6F |
| ASO6_PO_LNA | **CGoAoGoAoCAGTCGCTTCCoAoCTT** | 6F |
| ASO6_PS_cEt | **CGAGACAGTCGCTTCCACTT** | 6F |
| ASO6_PO_cEt | **CGoAoGoAoCAGTCGCTTCCoAoCTT** | 6F |
| ASO7 | **AAGGAGGTCATCCACGAAGT** | 2A-C; 6E; 7A-D |
| ASO7_PO | **AAoGoGoAoGGTCATCCACGoAoAGT** | 6E |
| ASO8 | **TGCATGGTGTAGCCCCCCTG** | 2A-C; 6E; 7A-D |
| ASO8_PO | **TGoCoAoToGGTGTAGCCCCoCoCTG** | 6E |

PO = **o** ; orange = MOE; black = DNA; blue = cEt;
red = LNA; *all linkages are PS unless noted

### Supplemental Table 2

**ASOs used in *in vitro* studies (1)**

| **ASO #** | **SimpleSequence** | **Figure #** |
|---|---|---|
| ASO2_PO | **GToToToToCAAACACACCToToCAT** | 9B |
| ASO3_PO | **GToToToToCTTATTTTTAGoCoATT** | 9B |
| ASO4_PO | **CAoCoToToCGAACTCCTGGoCoGGG** | 9B |
| ASO7_PO | **AAoGoGoAoGGTCATCCACGoAoAGT** | 9B |
| ASO8_PO | **TGoCoAoToGGTGTAGCCCCoCoCTG** | 9B |
| ASO35_LNA | **ATTTCCAAATTCACTT** | 9B |
| ASO36_LNA | **CTTTATTTCCAAATTCACTT** | 9B |
| ASO37_LNA | **ATTTCCAAATTCACTTTTAC** | 9B |
| ASO38_LNA | **TAGCCCTAAAGTCCCA** | 9B |
| ASO39_LNA | **CATGATTGTGGGCTTA** | 9B |
| ASO40_LNA | **ACTGGTTAGCCCTAAA** | 9B |
| ASO41_LNA | **CTTTATTTCCAAATTCACTT** | 9B |
| ASO35_cEt | **ATTTCCAAATTCACTT** | 9A-B |
| ASO36_cEt | **CTTTATTTCCAAATTCACTT** | 9B |
| ASO37_cEt | **ATTTCCAAATTCACTTTTAC** | 9B |
| ASO38_cEt | **TAGCCCTAAAGTCCCA** | 9A-B |
| ASO39_cEt | **CATGATTGTGGGCTTA** | 9B; 10A-B, |
| ASO40_cEt | **ACTGGTTAGCCCTAAA** | 9B; 10A-B, 10D |
| ASO41_cEt | **CTTTATTTCCAAATTCACTT** | 9B |
| ASO42 | **GCAAACAGGATACAGT** | 9A-B; 10A-B, 10D |
| ASO43 | **CTAGCCCACCCATCAATTTG** | 9B |
| ASO44 | **GTTAGCCCTAAAGTCCCAGG** | 9B |
| ASO45 | **TACATGCGTCCTTT** | 9B |
| ASO46 | **TTTATTTCCAAATTCACTTT** | 9B |
| ASO47 | **AAGATGAAATTTGCTC** | 9B |
| ASO48 | **TACTAGCCCACCCATC** | 9B; 10A, 10D |
| ASO49 | **AAAGATGAAATTTGCTCTTA** | 9B |
| ASO50 | **CCTTAATTTCACCCTC** | 9B; 10A, 10D |
| ASO51 | **CTTTATTTCCAAATTCACTT** | 9B |
| ASO52 | **GAoCoTAATATGCAGToTT** | 9B |
| ASO53 | **TToGoCCAATATCACCoAT** | 9B |
| ASO54 | **TToGoCCAATATCACCoAT** | 9B |
| ASO55 | **CAoAoCUGAACCACCCoGT** | 9B |

PO = **o** ; orange = MOE; black = DNA; blue = cEt; red = LNA; *all linkages are PS unless noted

### Supplemental Table 3

**ASOs used in *in vitro* studies (2)**

| ASO # | SimpleSequence | Figure # |
|---|---|---|
| ASO2_PO | GToToToToCAAACACACCToToCAT | 9C; 11A-E |
| ASO3_PO | GToToToToCTTATTTTTAGoCoATT | 9C |
| ASO4_PO | CAoCoToToCGAACTCCTGGoCoGGG | 9C |
| ASO5_PO | GCoCoAoGoGCTGGTTATGAoCoTCA | 9C |
| ASO6 | CGAGACAGTCGCTTCCACTT | 9C |
| ASO7_PO | AAoGoGoAoGGTCATCCACGoAoAGT | 9C; 11A-E |
| ASO8_PO | TGoCoAoToGGTGTAGCCCCoCoCTG | 9C; 11A-E |
| ASO9 | GAoToAoToTATCCTTTGAGoCoCAC | 9C |
| ASO10 | CCoGoToTTTCTTACCAoCoCCT | 9C |
| ASO11 | CCoToAoTAGGACTATCCAoGoGoAA | 9C |
| ASO12 | CAoGoAoCTGTAATCTAGGoAoCCC | 9C |
| ASO13 | ACoGoAoCATTTTCTTGCCoToCTT | 9C |
| ASO14 | GCoToCoATATCTAAAGACoCoGCA | 9C |
| ASO15 | TGoAATTCCTTTACACCAoCAC | 9C |
| ASO16 | TToTTTGTTAATAGTTCToCTG | 9C |
| ASO17 | CGoGoToAoAACTTTATATGoGoCTC | 9C |
| ASO18 | CAoGACTGTAATCTAGGoAoCCC | 9C |
| ASO19 | TCTCTATTGCACATTCCAAG | 9C |
| ASO20 | GCoToToTTACTGACCAoToGCG | 9C |
| ASO21 | CToCoToTACTCCCATCoAoCTG | 9C |
| ASO22 | TCoToGoTCTTTGGAGCoCoCAA | 9C |
| ASO23 | GCoToGoCGATCCCCAToToCCA | 9C |
| ASO24 | AAoToAoCATCCATGGCoTAA | 9C |
| ASO25 | GGoCoCoTTTGAAAGTCoCTT | 9C |
| ASO26 | TGoToAoTTTTGGATGCoTTC | 9C |
| ASO27 | TCoCTTTACACCACAoCTG | 9C |
| ASO28 | TToToTGTTAATAGTToCTC | 9C |
| ASO29 | TToToTGTTAATAGTToCTC | 9C |
| ASO30 | AToTCCTTTACACCACAoCoTGG | 9C |
| ASO31 | TToTTTGTTAATAGTTCoToCTG | 9C |
| ASO32 | GCoCoTTACTCTAGGAoCoCoAoAGA | 9C |
| ASO33 | ACoCoCoTTTCCATGTGAAoCoATT | 9C |
| ASO34 | AGoCoAoATCATTGGTAGCoAoTAC | 9C |

PO = o ; orange = MOE; black = DNA; blue = cEt; red = LNA; *all linkages are PS unless noted
